# Area use and residency behaviour at reproductive and foraging areas of green turtles from the largest rookery in the Atlantic

**DOI:** 10.64898/2026.09.02.749028

**Authors:** Jaime Restrepo, Daniel R. Evans, Raúl García-Varela, Angela Liu, Sandra Neubert, Kelor E. Cordero-Umaña, Lily K. Bentley, Anthony J. Richardson, Jeffrey A. Seminoff, Roldán A. Valverde, Daniel C. Dunn

**Affiliations:** Centre for Biological Conservation Sciences, The University of Queensland, St Lucia, QLD, Australia; Sea Turtle Conservancy, Gainesville, FL, United States; Centre for Ecology and Conservation, Faculty of Environment, Science and Economy, University of Exeter, Penryn, Cornwall, TR10 9FE, UK; Universidad Internacional Menéndez Pelayo (UIMP-CSIC), Madrid, Spain; The Leatherback Trust, Goldring-Gund Marine Biology Station, Playa Grande, Costa Rica; Commonwealth Scientific and Industrial Research Organisation (CSIRO) Environment, BioSciences Precinct (QBP), St Lucia, QLD, Australia; Marine Mammal and Turtle Division, NOAA Southwest Fisheries Science Center, La Jolla, California, USA; School of Earth, Environmental, and Marine Sciences, The University of Texas Rio Grande Valley, Brownsville, TX, USA

**Author notes:** Correspondence /.

**Keywords:** Green Turtles, Internesting movement, Foraging, Area Use, Marine Protected Areas coverage, Residency Index Utilisation Distributions

## Abstract

The conservation of migratory marine megavertebrates depends on the protection of different areas during their life cycles. Many populations have recovered following exploitation-driven declines. Green turtles exemplify successful efforts protecting reproductive populations, reducing harvesting, promoting population recoveries globally. Tortuguero, Costa Rica, the second largest rookeries for green turtles worldwide, implemented long-term conservation efforts leading to a population recovery, which is now in decline. This study investigates migratory connectivity of green turtles from Tortuguero across the Caribbean, and the role marine protected areas (MPAs) play in protecting key habitats supporting this population. Over 25 years, we satellite tagged 63 female green turtles to study their internesting and post-nesting foraging movements. Applying a state-space model, we calculated a movement-persistence index, to separate internesting and foraging periods. We calculated turtles’ utilization distribution areas for both periods. To assess the protection coverage at each end of the migration, we calculated time spent within and outside MPAs. Tracked turtles spent a median of 42.3 days (SD=18.8, range=12-80) within 30 km offshore before departing the internesting area, spending just 23.0% of time within nearby MPAs. After nesting, turtles migrated to 14 different foraging areas, Miskitos Cays, Nicaragua hosted the most turtles (n=23). Mean transmission time in foraging areas was 175.6 days (SD=126.0; range=13–544). Generally, turtles foraged 58.0% of the time within MPAs boundaries; however, ∼32% of tracked turtles foraged outside MPAs. Considering all known stressors, we found this population to be insufficiently protected at either end of their migratory cycle, remaining vulnerable to regional threats.

**Highlights:**

- Nesting green turtles at Tortuguero, Costa Rica, spend ∼20% of their internesting time within marine protected areas.
- We described area-use by green turtles in 11 foraging grounds never defined for this population; and confirm known connectivity between Tortuguero and at least 9 different countries, supporting previous work.
- Foraging area coverage by MPAs was relatively evenly split in thirds, with one third of foraging areas being entirely within MPAs, one third overlapping MPAs to some degree, and one third existing entirely outside MPAs.
- Almost half of the foraging grounds identified for green turtles across the Caribbean Sea and the southern Gulf of Mexico are outside MPAs, making turtles in these areas vulnerable to direct and indirect risks.

## 1. Introduction

Over the past century, many emblematic marine species have shown dramatic recoveries from earlier declines driven by exploitation or habitat degradation (Magera et al., 2013). The northern elephant seal (*Mirounga angustirostris*) has bounced back from near extinction driven by commercial sealing (Lowry et al., 2014). Similarly, numbers of humpback whales (*Megaptera novaeangliae*) have grown rapidly since their legal protection in 1955 (Cooke, 2018; Duarte et al., 2020), particularly the North Atlantic (Stevick et al., 2003) and Eastern Australian (Bejder et al., 2016; Noad et al., 2019). Increasing trends in marine turtle nesting populations in different regions suggest similarly positive conservation stories (Hays et al., 2025; Mazaris et al., 2017). Notably, the historically threatened green turtle (*Chelonia mydas*) was recently globally reclassified by the International Union for Conservation of Nature (IUCN) from “Endangered” to “Least Concern”, showing that protection of nesting females and their eggs, and reduction in harvest pressure, has led to population recovery in many places around the world (Wallace & Broderick, 2025). Nonetheless, some sea turtle populations are still declining and remain threatened by human activities and exploitation (Allen et al., 2023; Mancini et al., 2019; Restrepo et al., 2023).

The population of green turtles nesting in Tortuguero, on the Caribbean coast of Costa Rica, is the largest nesting assemblage of this species in the Atlantic Basin (Lahanas et al., 1998), and for many years was considered a prime example of successful marine migratory species conservation (Bjorndal et al., 1999; Troëng & Rankin, 2005). The nesting ecology of this population has been monitored for almost seven decades, making this one of the longest wildlife conservation programs worldwide (Restrepo et al., 2023). Prior to conservation efforts in Tortuguero, a large proportion of female green turtles encountered while nesting were harvested and sold to local and international markets (Carr, 1954). This intensive exploitation presumably drove the nesting declines observed in the first half of last century.

After a long-term collaborative intervention by the Costa Rican government (Government of Costa Rica, 1963; 1969; 1970; 2002) leveraging international treaties such as the Endangered Species Act (Endangered Species Act, 1973) and the Convention on International Trade in Endangered Species of Wild Fauna and Flora (CITES, 1973), the local nesting population showed a significant recovery (Figure 1; Gutiérrez-Lince et al., 2021) lasting nearly forty years (Bjorndal et al., 1999; Troëng & Rankin, 2005). However, in 2008, the annual estimated nest numbers of the Tortuguero green turtle population began a rapid decline, which continues today (Macias Nieto et al., 2025; Restrepo et al., 2023). The causes for this decline are not yet completely understood and raise questions about the longevity and sustainability of other conservation efforts for marine migratory species.

**Figure 1.**
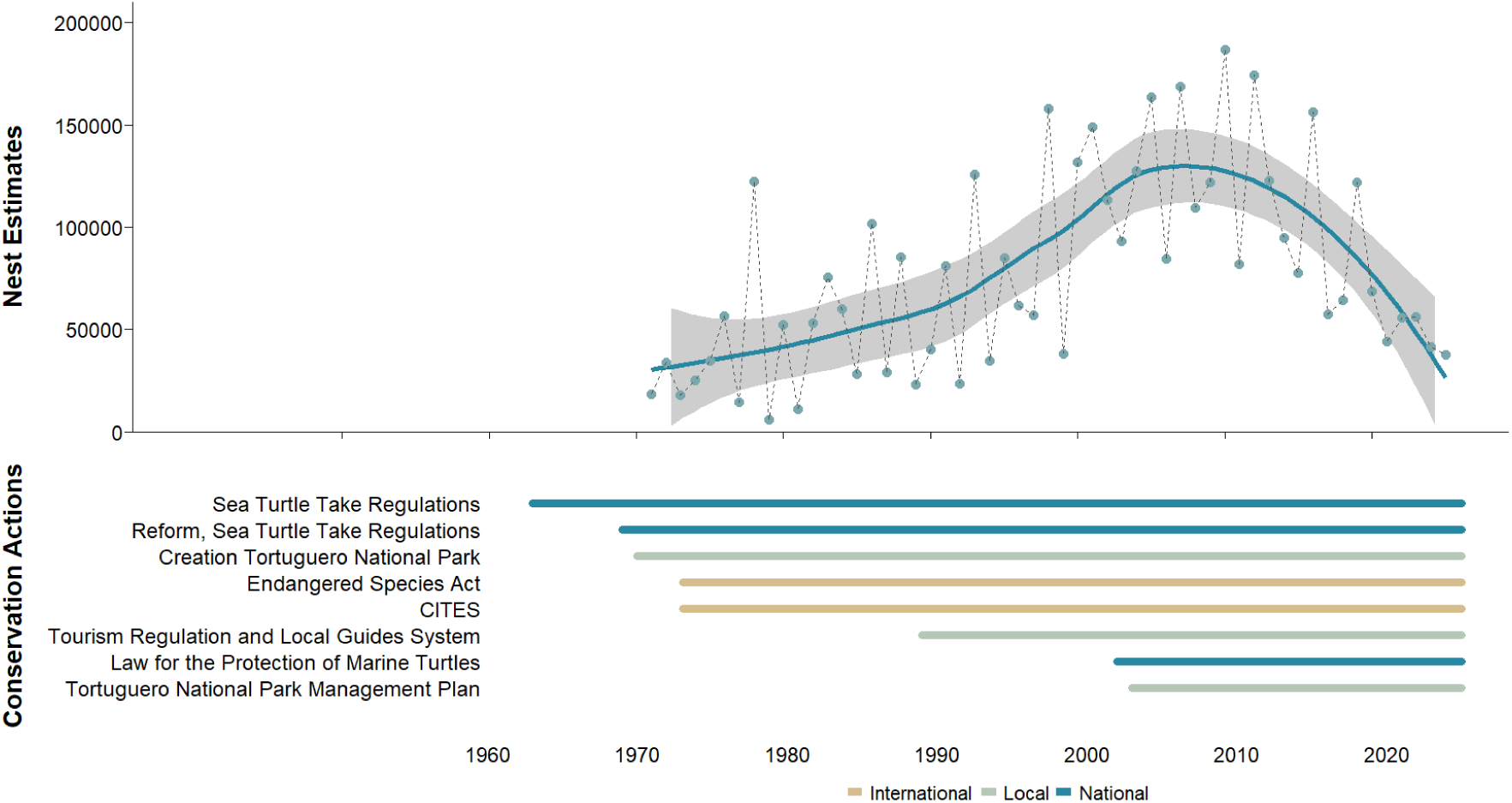
Nesting trend of green turtle (*Chelonia mydas*) (1971 – 2025) at Tortuguero, Costa Rica *(Macias Nieto et al., 2025; Restrepo et al., 2023)* and national, local and international conservation actions supporting marine turtle conservation.

Gaining a deeper understanding of key habitats could provide vital insights into why this specific nesting population is in decline. The persistence of many migratory marine megavertebrates – including many cetaceans, seabirds, sharks, rays and sea turtles – depends on access to key habitats across different life-history stages (Duarte et al., 2020; Lascelles et al., 2014). Migratory connectivity describes the cyclical movement of migratory individuals or populations between reproductive and foraging habitats (Marra et al., 2006; Webster et al., 2002). These key habitats are geographically dispersed, sometimes located tens of thousands of kilometres apart. Further, the ability of populations to access these habitats and their resources can have a profound influence on species ecology and evolution (Baguette & Van Dyck, 2007; Kool et al., 2013; Virtanen et al., 2020; Webster et al., 2002). Persistent threats in one habitat occupied during the life cycle of a migratory species can affect the status of the entire population (Runge et al., 2015). Thus, the conservation of migratory marine megafauna requires knowledge of the location, movement and habitat use of animals throughout the annual cycle (Pearson et al., 2020). Without information about the migratory connectivity of the population, managers are blind to the impacts that may be occurring outside of their jurisdiction, which may be undermining their conservation efforts.

Flipper tag recovery data from 50 years of tagging efforts at Tortuguero (Carr, et al., 1978) provides clear evidence that females embark on long migratory movements that span nearly the entire Caribbean Sea. Within this vast region, tag recoveries reveal the Miskito region of Nicaragua as the primary foraging ground for this population (Troëng et al., 2005).

Traditionally, Miskito Indigenous fishers have been permitted to legally harvest up to 12,000 green turtles annually (Lagueux et al., 2014). While this represents a pressure on the Tortuguero green turtle population, harvest in Nicaragua occurred throughout the apparent population recovery – indicating something else may be playing a role in declines (Bjorndal et al., 1999; Troëng & Rankin, 2005). Further, the true importance of this region as a foraging site for Tortuguero green turtles is not entirely clear, as tag returns from the Miskito region may have higher likelihood than those from other regions due to the encouragement of a monetary reward scheme that does not exist elsewhere (Troëng et al., 2005).

Satellite telemetry is a more accurate tool than mark-recapture methods in the assessment of marine turtle abundance and movement (Esteban et al., 2017; Tucker, 2010; Weber et al., 2013). Satellite tracking offers an independent and largely unbiased approach for quantifying movement patterns and habitat use, particularly when sample size is large enough; this technology has transformed our understanding of marine turtle spatial ecology and migratory behaviour (Bentley et al., 2025; Restrepo et al., 2026). By revealing habitat use, migratory routes, and connectivity among geographically separated habitats, telemetry can provide critical information for conservation planning and management (Wallace et al., 2010). Such information is particularly valuable for assessing cumulative impacts and the level of protection during different life stages (Gilmour et al., 2025; Trevail et al., 2025).

Marine protected areas (MPAs), when appropriately designated and managed, are one of the main management tools employed to protect threatened species and conserve biodiversity worldwide (Sanabria-Fernandez et al., 2019). MPAs are area-based management tools, aiming at preserving the health, productivity, and resilience of species assemblages and their supporting habitats (Grorud-Colvert et al., 2021). Effective management of marine ecosystems in the Caribbean Sea depends on the design, implementation and monitoring of conservation actions, especially MPAs (Cramer et al., 2026a). Nonetheless, despite their global importance, and a growing international interest in creating large-scale MPAs, the Greater Caribbean region has lagged behind in their implementation (Gallagher et al., 2020).

Here we show tracking data for 63 female turtles breeding at Tortuguero collected across 25 years and assess the population’s area-use and migratory connectivity to better understand the protection coverage from MPAs throughout its breeding cycle. We describe internesting and foraging habitats used by female green turtles throughout their reproductive migrations and assess protected area coverage at both ends of their migration. We discuss the implications of these insights for population migratory connectivity and provide an updated assessment of the relative importance of the Miskito foraging areas.

## 2. Methods

### 2.1. Tag deployment

Tortuguero National Park (TNP), on the northern Caribbean coast of Costa Rica, includes a ∼30-km stretch of beach that hosts the largest nesting assemblage for green turtles in the western hemisphere (Restrepo et al., 2023). This nesting population has been monitored via capture-mark-recapture efforts for seven decades. Further, between 2000 and 2024, 63 Argos platform transmitter terminals (PTTs) were attached on nesting green turtles. Most tagged turtles (n=41) were captured and released near Tortuguero Village (10.53779 °N, −83.50443 °W). The remaining 22 satellite transmitters were deployed between 2022 and 2024 on turtles nesting later in the season, on the beach sector known as “Jalova”, ∼20 km south of Tortuguero Village (10.38915 °N, −83.41396 °W; Table 1).

**Table 1.** Satellite tag deployments for nesting green turtles (*Chelonia mydas*) at Tortuguero, Costa Rica.

| *Nesting Season* | *N° Tags Deployed* | *Tag Brand* | *Tag type* | *Attachment Technique* | *Deployment Location* | *Reference* |
| --- | --- | --- | --- | --- | --- | --- |
| 2000 | 3 | Telonics | ST-14 | Fiberglass | Tortuguero | Troëng et al., 2005 + This study |
|  | 3 | Telonics | KiwiSat 101 | Fiberglass | Tortuguero | Troëng et al., 2005 + This study |
| 2001 | 3 | Telonics | ST-14 | Fiberglass | Tortuguero | Troëng et al., 2005 + This study |
| 2002 | 1 | Telonics | ST-14 | Fiberglass | Tortuguero | Troëng et al., 2005 + This study |
| 2009 | 1 | SirTrack | KiwiSat 101 | Epoxy (Simpson Strong-Tie) | Tortuguero | This study |
|  | 1 | Wildlife Computers | MK-10 GPS | Epoxy (Simpson Strong-Tie) | Tortuguero | This study |
| 2011 | 2 | SirTrack | KiwiSat 101 (K1G 291A) | Fiberglass | Tortuguero | This study |
| 2012 | 1 | SirTrack | KiwiSat 101 (K1G 291A) | Fiberglass | Tortuguero | This study |
| 2013 | 2 | SirTrack | KiwiSat 202 (K2G 575A) | Epoxy (Powers 308+) | Tortuguero | This study |
| 2014 | 2 | SirTrack | KiwiSat 202 (K2G 575A) | Epoxy (Powers 308+) | Tortuguero | This study |
| 2015 | 2 | SirTrack | KiwiSat 202 (K2G 576A) | Epoxy (Pure50+) | Tortuguero | This study |
| 2016 | 2 | SirTrack | KiwiSat 202 (K2G 576A) | Epoxy (Pure50+) | Tortuguero | This study |
| 2017 | 3 | SirTrack | KiwiSat 202 (K2G 576A) | Epoxy (Pure50+) | Tortuguero | This study |
| 2018 | 3 | SirTrack | KiwiSat 202 (K2G 576A) | Epoxy (Pure50+) | Tortuguero | This study |
| 2019 | 3 | SirTrack | KiwiSat 202 (K2G 576A) | Epoxy (Pure50+) | Tortuguero | This study |
| 2021 | 2 | Wildlife Computers | SPLASH10 | WC Attachment | Tortuguero | This study |
| 2022 | 2 | Wildlife Computers | SPOT-375B | WC Attachment | Tortuguero | This study |
|  | 6 | Wildlife Computers | SPOT-375B | WC Attachment | Jalova | This study |
| 2023 | 3 | Wildlife Computers | SPOT-375B | WC Attachment | Tortuguero | This study |
|  | 10 | Wildlife Computers | SPOT-375B | WC Attachment | Jalova | This study |
| 2024 | 2 | Wildlife Computers | SPOT-375B | WC Attachment | Tortuguero | This study |
|  | 6 | Wildlife Computers | SPOT-375B | WC Attachment | Jalova | This study |

### 2.2. Tag attachment

We intercepted turtles returning to the sea after oviposition, retaining them on the beach in wooden pens. For all transmitter-equipped turtles, we recorded or applied Inconel flipper tags (National Band and Tag Company, Style 681) to both fore flippers for individual identification. We then measured curved carapace length with a flexible tape measure to the nearest 0.1 cm (Wyneken, 2001), and conducted a visual inspection of their general body condition.

Prior to transmitter attachment, algae and epifauna were removed from the anterior portion of the carapace, the area around the second vertebral scute was roughened using 80-grit sandpaper, and the surface was cleaned with 70% ethanol to remove residues and natural lipids. As tag attachment practices evolved over the course of the study, satellite tags were attached using multiple techniques (Table 1) following protocols detailed in Evans et al. (2024).

### 2.3. Data processing

We collected 308,722 raw Argos locations from the 63 tracked green turtles. Data filtering first excluded any temporal or spatial duplicates, retaining a single fix per time and location. We then excluded all on-land locations, and filtered data by comparing 3 consecutive locations and removing unrealistic fixes indicating speeds > 9.9 km/h, and obtuse turning angles using the *SDLfilter* R package (Shimada et al., 2012).

We applied a state-space model to calculate a movement persistence index (ϒt) for every tracked turtle, based on their swimming speed and change in direction, using the *aniMotum* R package (Jonsen et al., 2023). High movement persistence is associated with migratory behaviour, when turtles travel in a linear trajectory at a relatively consistent, high speed.

Conversely, low movement persistence is indicative of slow movements and repetitive direction changes, interpreted either as inter-nesting movements (when low ϒ_t_ occurred at the beginning of a track) or residency in a foraging area after migration (when low ϒ_t_ occurred at the end of a track). These two behavioural periods of low move persistence were then separated out for further analysis.

### 2.4. Area-use models

For both internesting and foraging track segments, we calculated core and overall kernel utilization distributions (KUDs) and delineated the 50% and 95% probability contour areas for each tracked animal (hereafter UD50 and UD95) using the *adehabitatHR* R package (Calenge, 2023). UD95 provides unbiased home range estimates for the area occupied by turtles during each one of the behaviour time periods (internesting and foraging), even with relatively few data (Börger et al., 2006); whilst UD50 is the representation of core areas used by each turtle (Anderson, 1982).

To gather information on individual nesting turtles, we analysed female encounter records from 2015–2025. Based on flipper tag recapture data from nesting turtles (1,107 nesting encounters), we calculated the modal number of days between nesting events for green turtles re-encountered more than once within a single nesting season. We used this period to estimate internesting interval and the number of clutches laid by each turtle after tag deployment (Table S2).

Deployment date, transmitter malfunctions, and the potential take of turtles influenced the sample size for the analysis of each movement behaviour. Based on the combined locations for 45 tracked turtles remaining near the rookery during the internesting period, we evaluated the population-level internesting UD50 and UD95 around the nesting beach. Similarly, we determined the UD50 and UD95 for individual green turtles during each successive interesting interval between consecutive clutches.

Using the subset of tracking data collected post-migration, we calculated UD50 and UD95 for 56 foraging turtles. We grouped turtles in distinct foraging areas based on proximity and overlap of their individual foraging ranges. To be able to compare foraging areas independently of the individual foraging time, we assessed the individual weekly variation on UD50 proportioned to the area occupied by each turtle on the previous week (Figure S2).

### 2.5. Marine Protected Area coverage

We used the World Database on Protected Areas (WDPA), to identify MPAs used by tracked turtles. For each MPA, we extracted information on their IUCN conservation and management category (Day et al., 2019), size, and the year of establishment (UNEP-WCMC & IUCN, 2026; Table 3). To assess the occupancy of MPAs throughout both the internesting and foraging periods, we estimated a residency index (RI) by calculating the proportion of PTT locations transmitted by each turtle from within MPAs boundaries, over the total number of locations transmitted during the internesting/nesting or foraging period (Abalo-Morla et al., 2022; Revuelta et al., 2015). RI values within MPAs range from 0, indicating no time spent in MPAs, to 1, indicating constant residency within an MPA.

### 2.6. Space use and protection across different habitats

To understand space utilisation and the amount of time turtles spend in MPAs, we developed two statistical models for the internesting period at TNP, and two for the foraging period throughout the Caribbean Sea. All statistical models were fit using the *glmmTMB* R package (Brooks et al., 2017).

We first developed a generalised linear mixed model (GLMM) to investigate the relationship between variation in area of UD50 and the duration of the internesting period. Then we developed two GLMMs with different response variables: the Area (i.e. UD50) and the Residency Index (RI) for each internesting interval between nesting events. The two fixed predictors in each GLMM were the same: a factor for Clutch number, which varied from 1 to 6 for each turtle; and a factor for Beach Section, (Tortuguero/north or Jalova/south). We also included a random effect for TurtleID, as there were multiple estimates of Area or Residency Index for each turtle during the Internesting period (i.e., an Area and Residency Index was estimated during the time between each Clutch). For the internesting area, we first tried a normal error structure, but the diagnostic plots showed strong violations of the assumptions for the normality and homogeneity of variance, and we found that a gamma error structure with a log link function substantially improved the assumptions (Figure S3). For the internesting Residency Index, we used a beta error structure with a logit link function.

Because the Residency Index varies from 0 to 1, we first tried a GLMM using the ordbeta family (a beta error structure for data that contains 0s and 1s) but found that the model did not fit. We thus used a standard beta error structure with a “lemon squeezer”, which converts 0 to very small positive values and 1s to values just below 1 (Smithson & Verkuilen, 2006; see Supp materials for diagnostic plots).

#### Internesting model 1

Overall Area: glmmTMB(Total_Area_UD50 ∼ Nesting_Period, family = Gamma(link = “log”))

#### Internesting model 2

Clutch Area: glmmTMB(Area ∼ Clutch + Release Site + (1|TurtleID), family = Gamma(link = “log”))

#### Internesting model 3

Residency Index: glmmTMB(Res Index ∼ Clutch + Release Site + (1|TurtleID), family = beta_family(link = “logit”))

For the GLMs in the foraging period, we included two predictors: a fixed factor for Site for the 14 foraging areas the turtles travelled to; and a continuous variable for Time, which represented the length of fime that the turtles were tracked at each Site. No random effect for turtle was included because each observation represented an individual turtle. Similar to the internesting period, the response for one GLM was the foraging area, and the response for the second GLM was the foraging Residency Index. The same error structures were used as for the Area and Residency Index in the internesting period GLMMs (see the Supp for diagnostic plots).

#### Foraging model 1

Foraging Area: glm(Area ∼ Foraging Site + Time, family = Gamma(link = “log”))

#### Foraging model 2

Foraging Residency Index: glm(Res Index ∼ Foraging Site + Area, family =beta_family(link = “logit”))

## 3. Results

Out of the 63 nesting green turtles equipped with satellite transmitters, only one turtle failed to transmit successfully for more than three days and was eliminated from the remaining analysis (Table S1). The remaining turtles were tracked for a mean of 212.3 d (*n* = 62, SD = 126.9 d, range = 52–643 d). Transmission duration was independent of transmitter brand, unit specification and attachment technique (Table S1). From these 62 turtles, 17 departed immediately after tag deployment, thus internesting behaviour was analysed for 45 individuals. Similarly, some turtles stopped transmitting during migration, so 56 turtles that displayed a defined feeding area were included in the foraging analyses.

### 3.1. Internesting area

For the 45 tracked turtles that remained in the internesting area after deployment of satellite transmitters, median internesting period was 42.3 d (*n* = 45, SD = 18.8 d, range = 12-80 d). The combined core internesting area (UD50) for turtles that nested at least once after release (n = 45) was 158.92 km^2^. This area extended 40 km along the coast between the river mouth of Tortuguero river (4 km north from Tortuguero village), and Parismina. Tracked turtles remained near the deployment location on the north side of Tortuguero National Park. The internesting UD95 covered 1861.90 km^2^ and extended from the Costa Rica–Nicaragua border 45 km north of Tortuguero, to the Port of Limón 80 km to the south, with some disconnected areas extending over the border between Costa Rica and Panamá (Figure 2).

**Figure 2.**
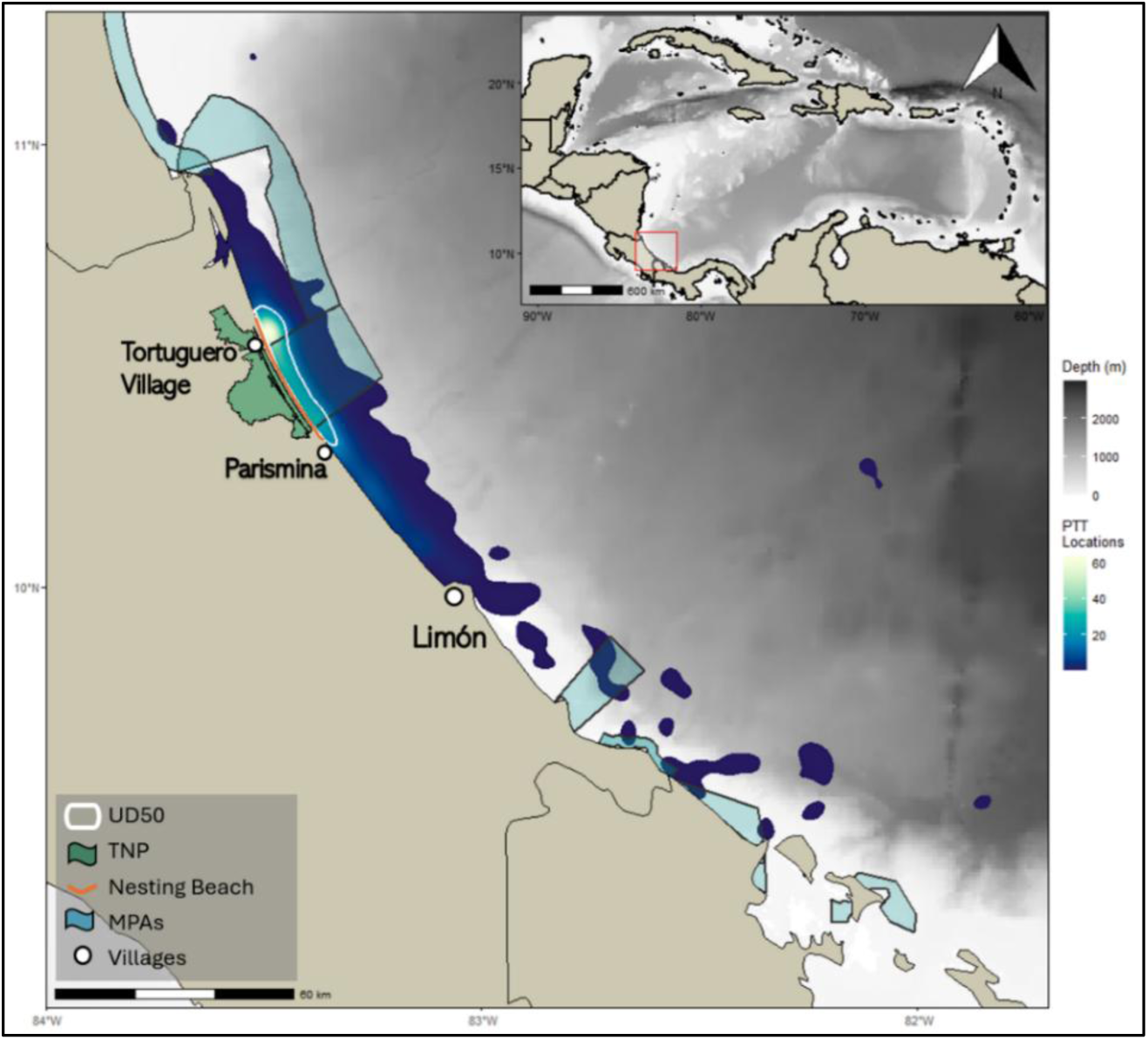
Map showing Kernel Utilization Distributions for the internesting period for green turtles (*Chelonia mydas*) in Tortuguero. The white contour represents the core internesting area (UD50), Blue shaded area represents home range area (UD95). The green polygon represents the land extension of Tortuguero National Park. White circles note the location of Tortuguero and Parismina villages as well as the commercial port of Limón. Light green polygons show the marine protected areas along the coast of Nicaragua, Costa Rica and Panamá *(UNEP-WCMC & IUCN, 2026)*. Blue-yellow scale shows the variation in density of PTT locations across the internesting area. Gray background shows bathymetry.

Individual internesting areas varied largely amongst all turtles, as UD50 area was not related to time spent nesting after tag deployment (−0.0035; SE=0.003; p-value=0.511; Figure 3; Table S4). All but two turtles (turtle id 19408 and 19585; <u>see supplementary figures</u>) remained <30 km from shore throughout the internesting period. These two individuals swam 160 km and 230 km away from the Tortuguero rookery, attending the Panamá-Colombia gyre between nesting events.

**Figure 3.**
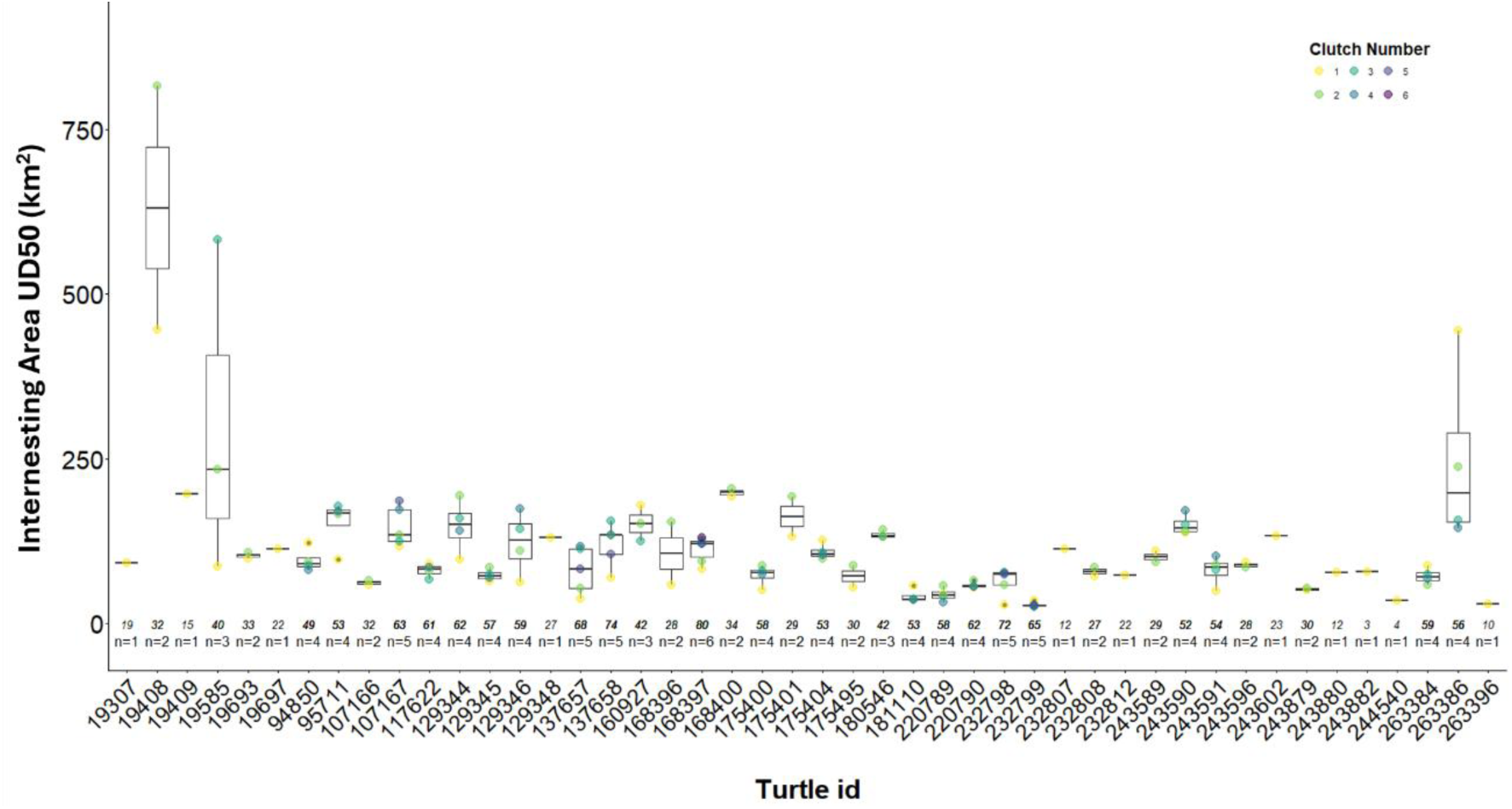
Internesting UD50 variation for individual nesting events of green turtles (*Chelonia mydas*) nesting at Tortuguero, Costa Rica. Boxplot showing the variance for UD50 throughout the individual nesting season of each tagged turtle. Data are presented chronologically by deployment date, numbers bellow the boxes represent the internesting period in days for each turtle, and the *n* values bellow that represents the number of estimated clutches laid by each individual.

Internesting interval was calculated to be 10 days between nesting events (mean = 30.3 d; sd = 18.34 d; range = 4–105 d; Figure S1). Internesting UD50 for single nesting events (i.e. 10 day periods) ranged from 25.40 km^2^ to 816.67 km^2^ (Table S2). These internesting intervals presented significant variations in UD50 between different nesting events over time (Figure 3). UD50 area for single nesting events increased throughout the nesting season (Figure 4a), and release location had a significant effect on UD50 area during the internesting period (Figure 4b; Table S4).

**Figure 4.**
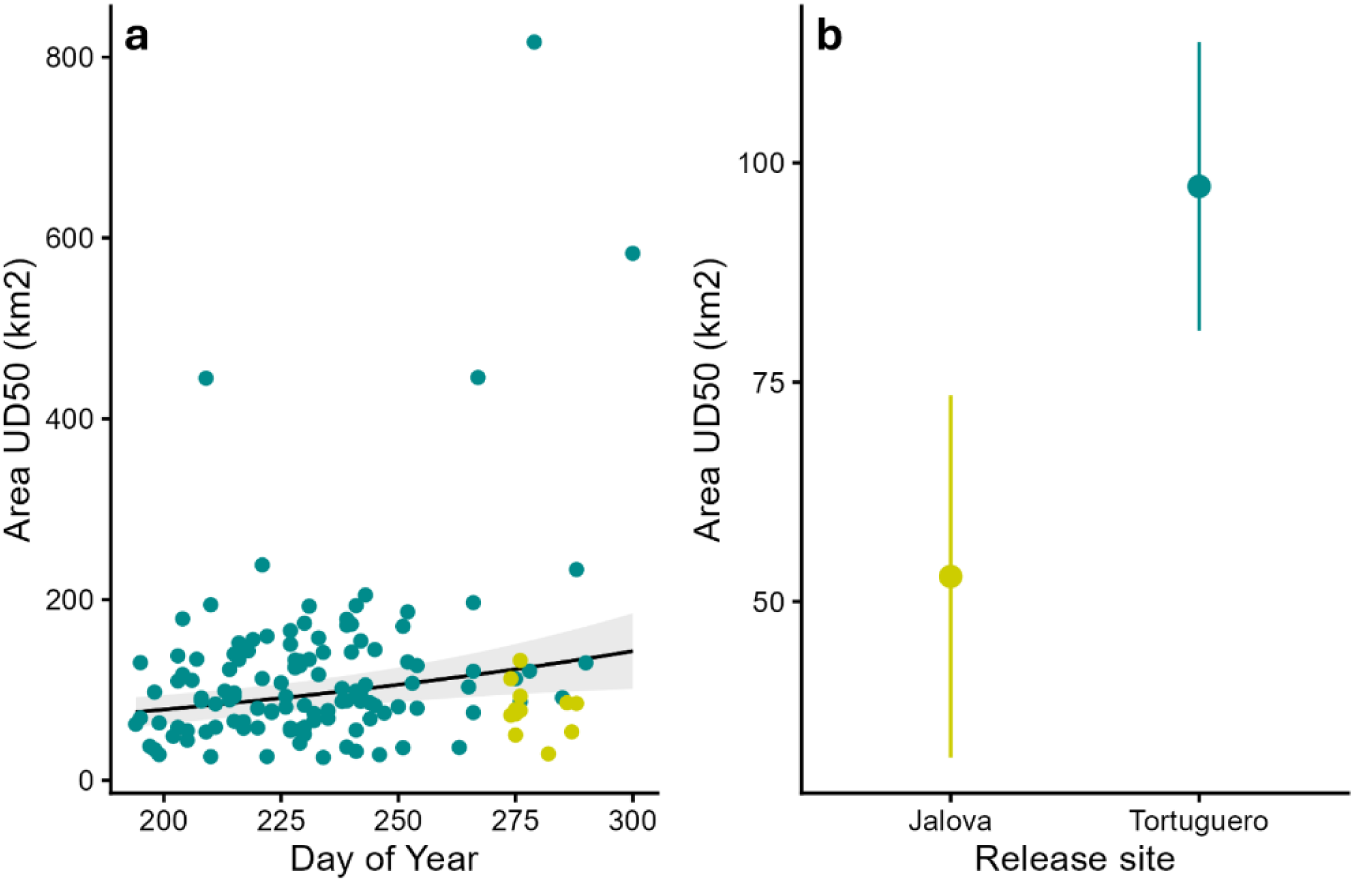
Predictions from the Generalised Linear Models during internesting period, assessing UD50 area variation for single nesting events of time for green turtles (*Chelonia mydas*) tracked from Tortuguero rookery (2000 - 2024). a) Area variation over time represented as day of the year. b) Area variation related to deployment location, “Tortuguero” (green) and “Jalova” (yellow).

### 3.2. Foraging

After concluding their individual nesting season at Tortuguero, each turtle returned to their respective foraging ground. After departure, green turtles were tracked to 14 distinct foraging grounds distributed across 9 countries in the Caribbean Sea (Figure 5), 52 turtles travelled directly north, four travelled east to foraging grounds in insular Caribbean, and one turtle travelled south to the Guajira in the northern Caribbean region of Colombia, where it seemed to have slowed down and initiated foraging behaviour. However, after only two days, we lost transmission. Due to the limited number of locations for the foraging behaviour of this turtle, we excluded this track from all analysis, since it was not possible to calculate foraging UDs. Of the 56 turtles that successfully arrived at foraging grounds, 50 reached foraging areas over the continental shelf and six attended different islands in the northeast Caribbean Sea (Figure 5). We found that continental foraging areas were in coastal (<60 km offshore) shallow waters (<50 m deep), and insular foraging areas were <35 km to land.

**Figure 5.**
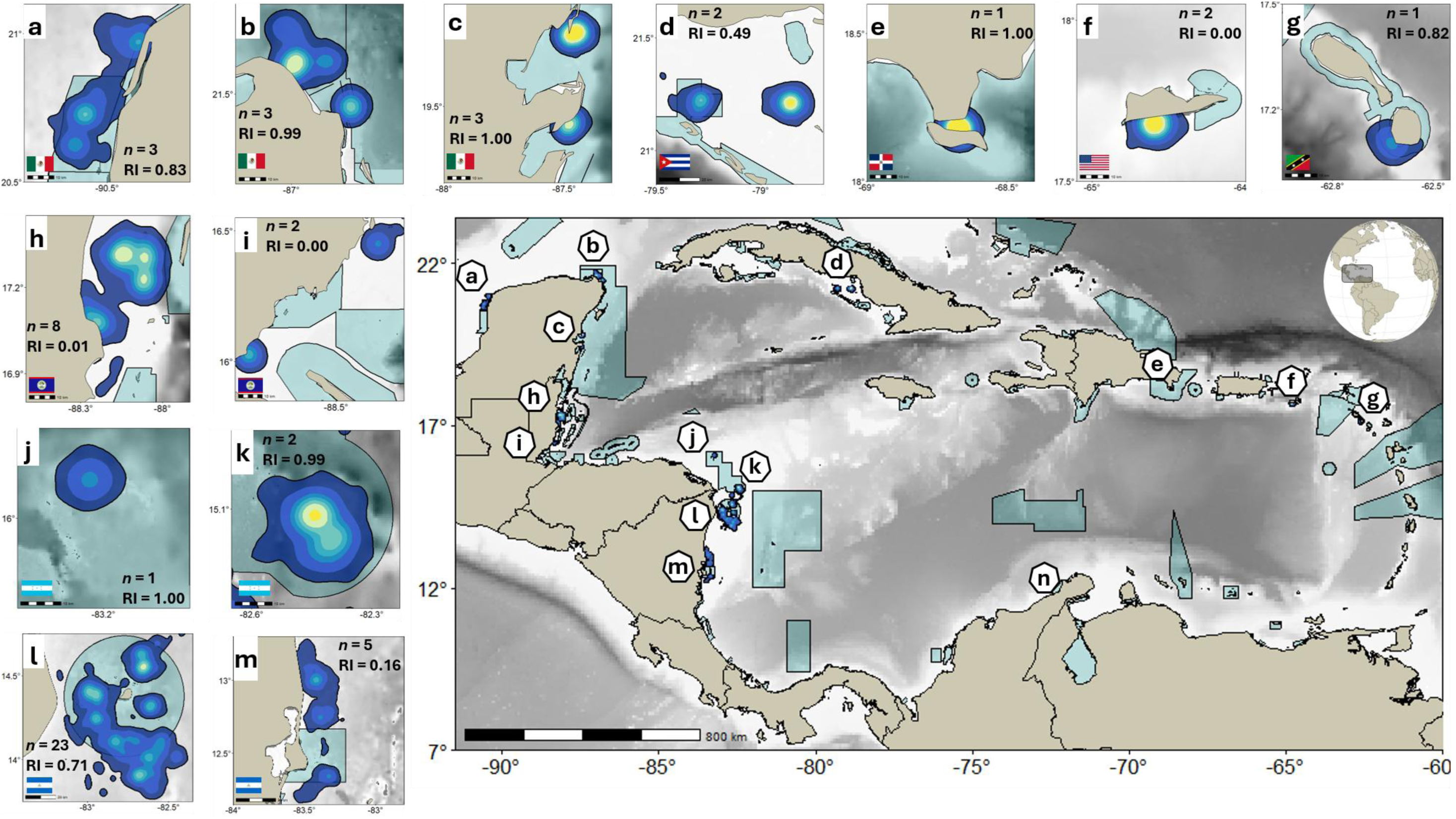
Foraging areas and kernel utilization distributions (KUD) for green turtle (*Chelonia mydas*) tracked from Tortuguero beach, Costa Rica from 2000 to 2024. Green shaded areas represent MPAs (UNEP-WCMC 2026); grey shading shows the bathymetry. KUD of foraging turtles at: a) Campeche, Mexico; b) Yucatán, Mexico; c) Quintana Roo, Mexico; d) Jardines de la Reina, Cuba; e) Sonoa Island, Dominican Republic; f) the US Virgin Islands; g) St. Kitts and Nevis; h) Belize; i) south Belize; j) Gracias a Dios, Honduras; k) Cayos Misquitos, Honduras; l) Miskitos Keys, Nicaragua; and m) Sandy Bay Sirpi, Nicaragua; n) Guajira, Colombia. *n* represents the number of turtles tracked to each one of the foraging grounds. RI is the MPA residency index.

The mean time animals were tracked in foraging areas was 172.8 d (*n*=56, SD=126.7 d, range=13–544 d), though this was routinely cut off by tag failure rather than return migration. All turtles remained in their foraging ground until tags stopped transmitting. Mean foraging UD50 was 55.5 km^2^ (*n*=56, SD=38.5 km^2^, range=20.6 – 237.1 km^2^; Table S2). Time foraging had a significant effect on the area size utilised by green turtles (Figure 6a; Table S4). Foraging UD50 differed widely among foraging grounds (Figure 6b), the largest being Miskitos and Sandy Bay Sirpi in Nicaragua (Table S4).

**Figure 6.**
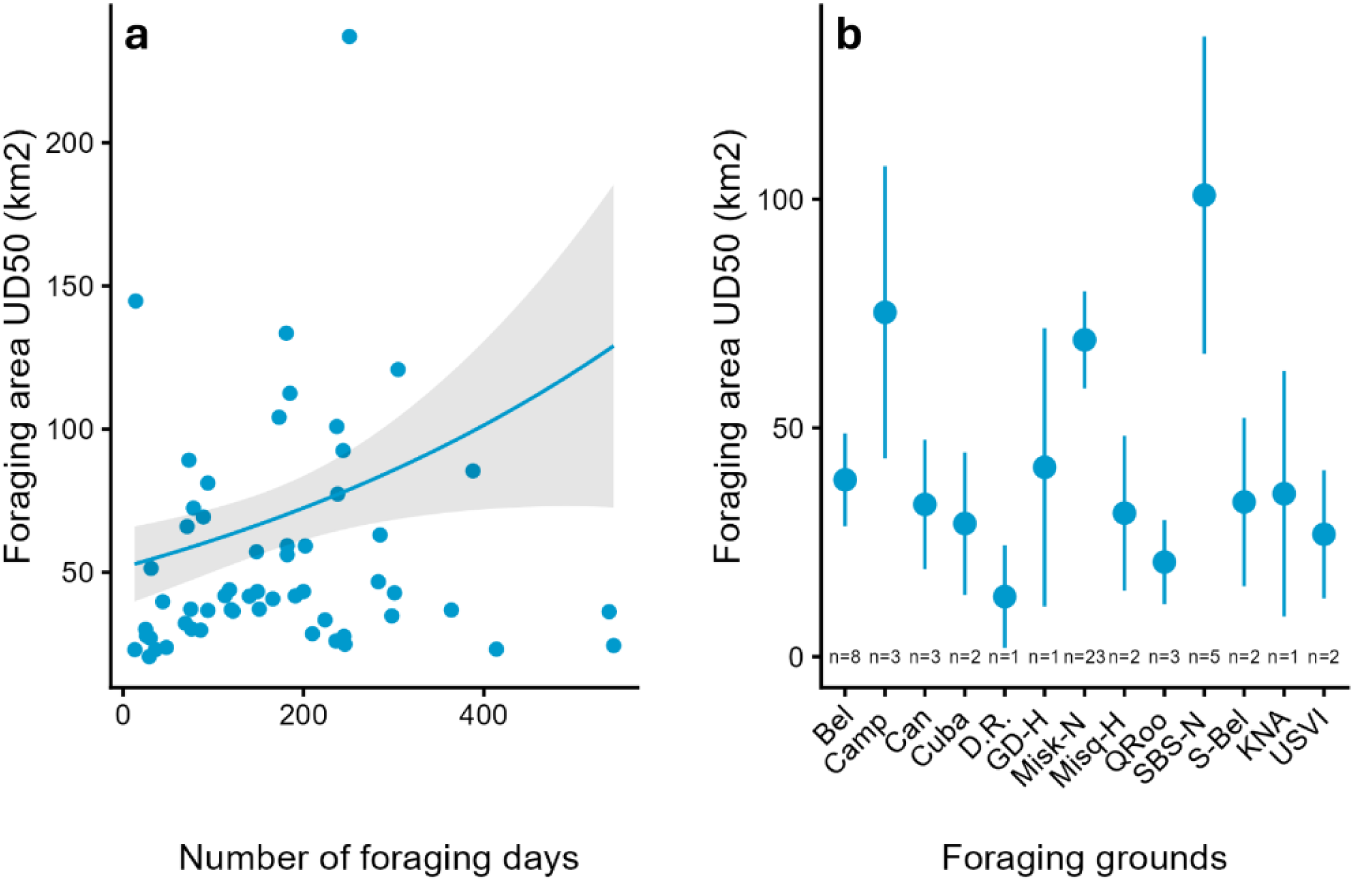
GLM predictions of UD50 area for green turtles (*Chelonia mydas*) tracked from Tortuguero rookery (2000 - 2024). a) UD50 variation over number of days spent foraging. b) UD50 variation for different foraging grounds. *n* = number of turtles tracked at each foraging ground.

### 3.3. Marine protected areas coverage

Turtles that remained at the rookery following tag deployment had an MPA RI of 0.22 (*n* = 2,022 locations in an MPA from 9,006 total locations). Individual RI within MPAs ranged from 0.04 to 1 (Table S2). MPA RI did not significantly vary across different internesting intervals (Figure 7a). Similarly, the number of clutches laid did not influence the time spent within MPAs (Table S4). Turtles released from Jalova had a significantly higher MPA RI than the those from Tortuguero (Figure 7b). MPAs attended during internesting were the fishing management area Barra del Colorado (3.0%), Cahuita National Park (5.6%), and Tortuguero National Park (89.9%) (Figure 5).

**Figure 7.**
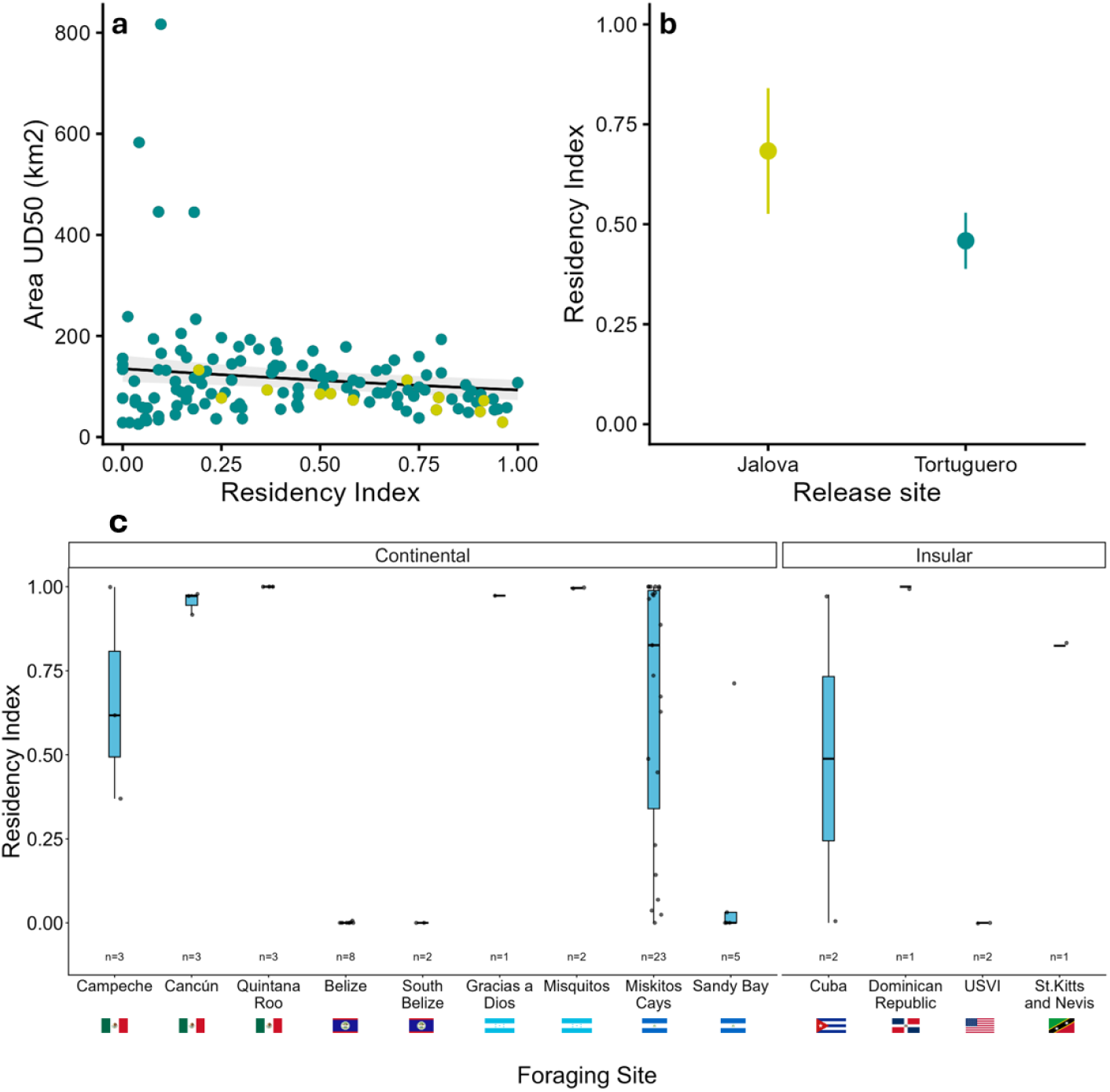
MPA Residency Index (RI) for internesting (top panel) and foraging (bottom panel) period for green turtles (*Chelonia mydas*) tracked from the Tortuguero nesting rookery (2000– 2024). a) Predictions from the Generalised Linear Models during internesting period, assessing the relationship between area used between consecutive nesting events and RI within MPAs around Tortuguero rookery. b) Predictions from the Generalised Linear Models during internesting period, comparing RI within MPAs for turtles released Jalova in yellow, and Tortuguero in green. c) Boxplot showing the RI within MPAs across foraging grounds in 8 different countries, considering continental and insular destinations; *n* = number of turtles within each foraging ground.

MPA RI during foraging varied, with about 17 of the tracked turtles totally inside MPAs (i.e. RI=1), 21 overlapping MPAs to some degree, and 18 not entering any MPAs. 46.4% of all foraging UD50s and 32.8% of foraging UD95s were entirely within the limits of MPAs, including some turtles feeding in Mexico, Miskito Keys, Nicaragua, Honduras, and Dominican Republic (Figure 5; Table 2). The 31% of turtles that did not enter the protection of any MPA while foraging included all turtles foraging in the territorial waters of Belize (n=10; Figure 5h,i); Nicaragua, Miskitos (n=1; Figure 5l) and Sandy Bay (n=3; Figure 5m); the U.S. Virgin Islands (n=2; Figure 5f); Campeche, Mexico (n=1; Figure 5a); and Cuba (n=1; Figure 5d). The 36.2% of the turtles that moved in and out MPA borders were in Miskito Keys and Sandy Bay, Nicaragua (n=16 and n=2, respectively; Figure 5l, m). Foraging grounds used by this population overlapped with 14 different MPAs, with varying levels of protection across the Caribbean Sea (Table 3). The mean foraging MPA RI was 0.53 (SD = 0.44; Table S2) and significant differences amongst foraging ground (Table S4; Figure 7c). Foraging grounds in Mexico, Honduras, Dominican Republic and St Kitts and Nevis had higher MPA Ris (>0.80), whereas Belize and the U.S. Virgin Islands had an MPA RI close to 0, except for a few detections inside the Turneffe Atolls Marine Reserve in Belize. MPA RIs within Nicaragua varied from 0.16 to 0.71 (Table 2).

**Table 2.** Characterization and use of foraging grounds for tracked green turtles (*Chelonia mydas*) nesting at Tortuguero, Costa Rica. Describing for each foraging ground: the Exclusive Economic Zone (EEZ), where country names are abbreviated MEX = Mexico, Bel = Belize, HND = Honduras, NIC = Nicaragua, CUB = Cuba, DOM, Dominican Republic, USA = United States of America, KNA = St. Kitts and Nevis, and COL = Colombia. Area occupied by the turtles is expressed as the mean and range of UD95. IUCN protection level *(UNEP-WCMC & IUCN, 2026)*. MPA Residency Index (RI) is an indicator of the time turtles spend inside MPAs.

| EEZ | Foraging Ground | N° Turtles | Mean UD95 (km<sup>2</sup>) | SD ± | Range UD95 (km<sup>2</sup>) | WDPA MPA Name* | Mean RI within MPAs | SD ± | RI within MPAs | Overall RI within MPAs | Figure |
| --- | --- | --- | --- | --- | --- | --- | --- | --- | --- | --- | --- |
| MEX | Campeche | 3 | 455.0 | 344.1 | 135.3 - 931.6 | Ría Celestún | 0.66 | 0.26 | 0.37 - 0.99 | 0.83 | Figure 5a |
|  |  |  |  |  |  | Los Petenes |  |  |  |  |  |
| MEX | Cancún | 3 | 184.4 | 48.2 | 117.0 - 226.5 | Tiburón Ballena | 0.96 | 0.03 | 0.92 - 0.98 | 0.99 | Figure 5b |
|  |  |  |  |  |  | Isla Contoy |  |  |  |  |  |
|  |  |  |  |  |  | Reserva de la Biosfera |  |  |  |  |  |
| MEX | Quintana Roo | 3 | 102.8 | 19.8 | 82.1 129.5 | Sian Ka ´ an | 1.00 | 0.00 | - | 1.00 | Figure 5c |
|  |  |  |  |  |  | Arrecifes de Sian Ka'an |  |  |  |  |  |
| CUB | Jardines de la Reina | 2 | 198.3 | 9.1 | 189.3 - 207.4 | Jardines de la Reina | 0.49 | 0.49 | 0.00 - 0.98 | 0.49 | Figure 5d |
| DOM | Sonoa | 1 | 46.8 | - | 46.8 | Cotubanamá (Del Este) | 1.00 | - | - | 1.00 | Figure 5e |
| USA | Virgin Islands | 2 | 81.5 | 6.8 | 74.7 - 88.4 | - | 0.00 | - | - | 0.00 | Figure 5f |
| KNA | St Kitts & Nevis | 1 | 107.0 | - | 107 | Saint Kitts and Nevis Marine Management Area | 0.82 | - | - | 0.82 | Figure 5g |
| BEL | Belize | 8 | 182.4 | 58.5 | 89.2 - 255.3 | Turneffe Atolls | 0.01 | 0.00 | 0.00 - 0.01 | 0.01 | Figure 5h |
| BEL | South Belize | 2 | 129.3 | 44.8 | 84.5- 174.1 | - | 0.00 | - | - | 0.00 | Figure 5i |
| HND | Gracias a Dios | 1 | 582.2 | - | 582.2 | Cayos Misquitos | 0.97 | - | - | 0.97 | Figure 5j |
| HND | Cayos Misquitos | 2 | 206.4 | 7.9 | 198.4 - 214.3 | Cayos Misquitos | 1.00 | 0.00 | 0.99 - 1.00 | 1.00 | Figure 5k |
| NIC | Miskitos Cays | 23 | 494.6 | 390.6 | 153.4 - 1957.7 | Cayos Miskitos y Franja Costera Inmediata | 0.66 | 0.38 | 0.00 - 1.00 | 0.71 | Figure 5l |
| NIC | Sandy Bay Sirpi | 5 | 525.0 | 232.6 | 199.9 - 843.5 | Cayos Perlas | 0.15 | 0.28 | 0.00 - 0.71 | 0.16 | Figure 5m |
| COL | Guajira | 1 | - | - | - | Pastos Marinos Sawairu | - | - | - | - | Figure 7n |

**Table 3.**
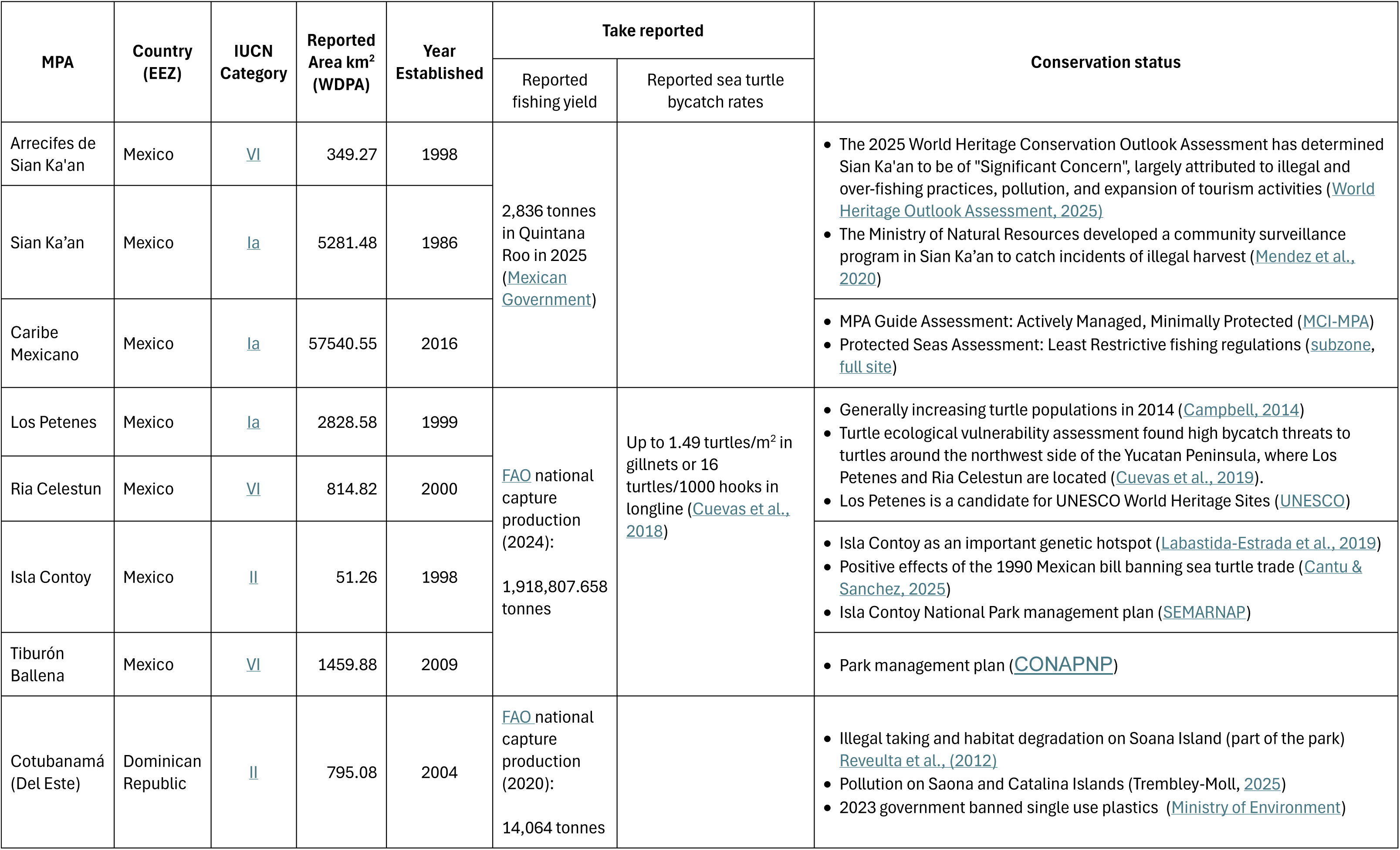
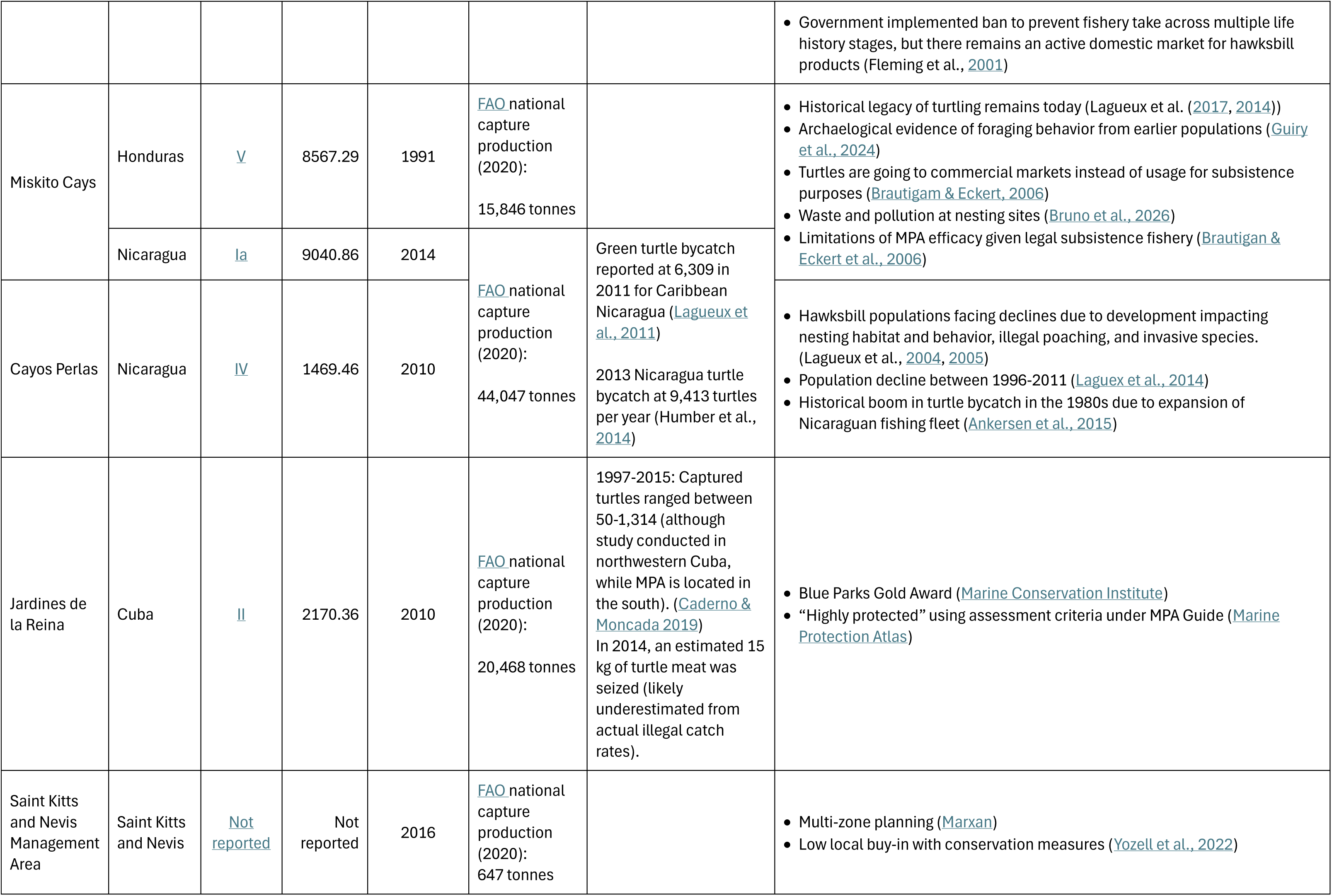
Characterization of Marine Protected Areas (MPAs) in different foraging grounds across the Caribbean Sea identify by tracked green turtles (Chelonia mydas) nesting in Tortuguero, Costa Rica (2000 – 2024). MPAs assessment criteria include country of jurisdiction; IUCN Protected Area Management Categories; Extension as the reported area in km^2^; Year of establishment; the take reported as legal, incidental or targeted anthropogenic interactions; and the conservation status accordingly to specific management plans, national assessments or independent research.

## 4. Discussion

Globally, the use of satellite telemetry has been a powerful tool for deciphering the complexities of animal ecology, including movement, habitat use and species interactions (Hussey et al., 2015). Our long-term tracking of green turtles from Tortuguero nesting beach expanded our understanding of their migratory connectivity and identified core use areas, both in internesting habitats and foraging grounds. We found that internesting movements were concentrated in shallow waters (<60 m) close to Tortuguero National Park (TNP), but turtles only spent ∼20% of their time within park boundaries. This low but consistent MPA residence during internesting was contrasted with more variable MPA coverage of foraging areas. Turtles were tracked to foraging areas in nine countries across the continental and insular Caribbean. While the Miskito region in Nicaragua was the most visited foraging area, most turtles dispersed to other foraging sites, many in coastal Belize and Mexico.

### 4.1. Large ranges and insufficient protection for nesting turtles

Internesting areas for green turtles at Tortuguero were generally larger than those reported in similar studies elsewhere (e.g. Goodwin, 2025; Mettler et al., 2020; Okuyama et al., 2025). We found that the internesting activity of tracked turtles was highly concentrated north of the limits of TNP, between Tortuguero Village and the river mouth (Figure 2). These turtles remained close to shore, often attending river mouths, where they may opportunistically feed during the nesting season, particularly on riparian vegetation brought downstream via runoff (Saragoça Bruno et al., 2026a). Generally, breeding green turtles display energy-saving strategies, frequently having resting dives in between nesting events (Hamann et al., 2002). However, when food is available within the internesting areas, green turtles will forage and supplement their energy budget during breeding season (Okuyama et al., 2025). The large areas used by green turtles during the nesting season may be related to their constant movement towards river mouths, expanding their range further than the nesting beach and outside the limits of Tortuguero National Park MPA.

The internesting MPA RI for this population shows that green turtles spend about 23% of their time within the limits of MPAs off the coast of Costa Rica, primarily in the marine area of TNP, which from June to August is used by this population as a courtship ground. This MPA coverage for tracked turtles is low, having nesting females spending just under a quarter of their internesting period within MPAs. Green turtles nesting at Tortuguero exhibit a distinctive high nest-site fidelity (Carr, & Carr, 1972; García-Varela & Harrison, 2016), we captured and released most of the tracked turtles in this study from the northernmost section of the nesting beach off limits of TNP, where according to Troëng and Rankin (2005), an estimate of between 1,600 and 3,300 green turtles nest every season. Thus, for the greater part of their internesting period, these females reside in unprotected coastal areas, where they are vulnerable to poaching (Rojas-Cañizales et al., 2022).

Tortuguero National Park was created with the main purpose of protecting the nesting population (Government of Costa Rica, 1970), and with the implementation of different conservation strategies over several decades, TNP has exhibited an increase in the activity of emblematic species (e.g. Guilder et al., 2015). However, based on our findings, the coverage of the Tortuguero MPA seems insufficient to protect the area used by green turtles during their interesting period. Further, despite all the long-term conservation efforts by Costa Rican authorities, poachers have adapted to law enforcement responses and have managed to continue the illegal take of nesting females and egg clutches from this rookery (Mejías-Balsalobre et al., 2021; Pheasey et al., 2021; Rojas-Cañizales et al., 2022).

We showed the nesting aggregation of green turtles at Tortuguero moving far beyond the limits of TNP, using over 115 km of coastline covering almost the entirety of the Caribbean coast of Costa Rica, but concentrating primarily north of the provincial capital of Puerto Limón (Figure 2). The commercial port of Puerto Limón is the 10^th^ largest port in Latin America, moving ∼13% of cargo volume relative to the Panama Canal. This heavy marine traffic, causing water pollution, acoustic disturbances and collision risk, impacting marine populations globally, limiting their natural movement (Barco et al., 2016; Pasanisi et al., 2022; Womersley et al., 2022). Green turtles in coastal areas have shown to unlikely take evasive actions to avoid collision with vessels traveling at speed greater than 4 km h^-1^ (Hazel et al., 2007), increasing the risk of collision and injury, particularly in turbid water. The port of Limón appears to be a limit for the continuous movement of green turtles nesting in Tortuguero since most turtles spent their internesting period swimming between this port and the nesting beach. Nonetheless, we identified at least four turtles moving farther south past the port (<u>see supplementary internesting figures</u>), two of these followed the Panamá-Colombia gyre and returned to the nesting beach, the other two crossed for a brief time to Cahuita National Park and across the border to the province of Bocas del Toro, Panamá, both known mating and foraging grounds for green turtles (Meylan et al., 2013; Moya-Ramírez et al., 2025).

### 4.2. Diffused foraging connectivity

Post-nesting migration of green turtles tracked from Tortuguero dispersed widely across the Caribbean Sea and the southeast of the Gulf of Mexico, confirming known connectivity based on metal tag returns data. Tracked turtles showed connectivity with at least 14 different, well-defined foraging areas where turtles established residency (showing no further displacement). Previous work based on flipper tag recaptures has shown that green turtles from Tortuguero have strong connections with foraging grounds in Nicaragua, Belize and Cuba (Carr, et al., 1978; Troëng et al., 2005). These tag recoveries also suggested a diffuse connectivity for this population, linking Tortuguero with at least 20 different foraging grounds across the Caribbean Sea. However, tag recaptures across the region probably does not reflect an unbiased distribution of connectivity for nesting turtles. Tag recapture information is normally obtained only along coastlines inhabited by fishermen and locations where specific research is conducted (Bjorndal et al., 2003; Richards et al., 2024; Troëng et al., 2005), ignoring isolated areas that may be important for the foraging ecology of this population. The relative importance of foraging destinations to the Tortuguero nesting population differs markedly depending on whether evidence came from satellite tagging or recovery of flipper tags (e.g. Richards et al., 2024).

Previous tracking studies on green turtles nesting at Tortuguero (including 10 of the turtles in this study; Table 1) showed 80% of tracked turtles foraging in shallow waters of Nicaragua and the remaining 20% going to Belize and Honduras (Troëng et al., 2005). Although the seagrass beds around the Miskito Cays in Nicaragua were still the main foraging area identified in this study, almost 60% of tracked green turtles travelled to other locations around the Caribbean Sea. The enhanced understanding of green turtles’ migratory patterns from Tortuguero provided by this study suggests a population with weaker migratory connectivity (i.e., a population that is more dispersed across diverse foraging areas) (Webster et al., 2002). This has implications for our understanding of the current population decline, in that it suggests that the population should be more resilient to impacts from artisanal fisheries in Nicaragua (Lagueux et al., 2014, 2017), forecasting more areas than previously thought (because a smaller proportion of the population is foraging there).

Conversely, it means that impacts on foraging grounds elsewhere have a higher likelihood of exacerbating the losses in Nicaragua and contributing meaningfully to the current population decline. We identified some foraging areas by the track of single turtles (i.e. St. Kitts and Nevis). Thus, we cannot be certain of the relative importance, or contribution of these areas to sustaining the nesting population at Tortuguero. Nonetheless, the identification and protection of distinct foraging grounds around the region may hold the key to the recovery of this declining nesting population, given the extended time these animals spend foraging in single locations, which may increase their vulnerability. This broader view of where turtles from Tortuguero are foraging, increases the interdependence of conservation strategies implemented across the region (Maina et al., 2020; Marra et al., 2006).

Foraging UDs calculated for the 56 tracked green turtles were all distributed in shallow-coastal areas, close to land (within 60 km of shore) either on the continental shelf or around small islands in the Caribbean. Some of the foraging grounds with higher number of tracked turtles (e.g. Miskitos Cays and Cayos Perlas; Figure 5), showed large ranges in UDs described by individual turtles (Table 2). These large variations may be due to a high density of foraging turtles competing to forage within a limited habitat (Hays et al., 2024). Sea grass meadows are not just important for supporting green turtle populations, they play a significant role in supporting fisheries through provision of nursery areas and trophic subsidies to adjacent habitats (Nordlund et al., 2018; Unsworth et al., 2019). Long-term studies in the Caribbean have shown important shifts in abundance, biodiversity and community structure of seagrass beds in the region (Fourqurean et al., 2019; Patriquin et al., 2024; van Tussenbroek et al., 2014). Continuous decline in abundance and range constrictions of seagrass beds in Bermuda have resulted in an increased competition for feeding resources, leading to reduction in body condition and early departure of immature green turtles from this important developmental area (Meylan, et al., 2022). These effects have probably cascaded over other important foraging grounds for green turtles around the Caribbean Sea, increasing the density of foraging turtles in already stressed ecosystems (Christianen et al., 2023; Heithaus et al., 2014). Further studies and comprehensive characterizations of foraging areas are needed to better address the conservation of seagrass meadows and the declining populations of marine turtles in the Caribbean Sea, and its impact on the nesting population at Tortuguero.

### 4.3. Marine protected area coverage variability across foraging areas

This study describes the area use patterns of green turtles foraging in at least 11 locations that have not been previously described for this population (Figure 5) and have different mixes of anthropogenic stressors that could contribute to population declines. We estimated the residency index for foraging green turtles to be highly variable with no protection for almost half of the foraging grounds across the Caribbean Sea. We identified at least six unprotected or partially protected foraging grounds used by green turtles in different countries around the Caribbean (Table 2). Making turtles in these areas vulnerable to direct stressors, such as boat strikes, entanglements, or direct take, as well as the indirect effects of noise pollution and water contamination may be encountered by green turtles in unprotected foraging grounds (Finlayson et al., 2016; Lagueux et al., 2017; Senko et al., 2022; Wallace et al., 2013).

The Yucatán peninsula in Mexico hosted nine turtles in three different areas with high coverage by MPAs. The use of seagrass meadows around the Yucatán peninsula has also been identified as an important area for other nesting populations of green turtles in the Caribbean (Cuevas et al., 2022; Uribe-Martínez et al., 2021). Further, in a comprehensive study, Cuevas et al. (2018) identified the potential impact of longline and gillnet fisheries on sea turtles around the Yucatan Peninsula, defining potential bycatch hotspots. Belize has the second most turtles foraging in its waters after Nicaragua, exploitation and consumption of marine turtles has also been previously reported in Belize (Moll, 1985). Even within the limits of MPAs, marine turtle populations in Belize are susceptible to several threats, primarily from anthropogenic origin (Delgado, 2018). Similarly, the remaining tracked turtles went to shallow areas in Colombia, Cuba, Dominican Republic, St. Kitts and Nevis, and U.S. Virgin Islands, places where marine turtles have traditionally been exploited in the past (Fleming, 2001). Even though, illegal take of marine turtles has decreased ∼60% in the past decades in the Caribbean (Humber et al., 2014), this region is still considered as a “high risk” and “high exploitation” for green turtles (Senko et al., 2022). Addressing specific regional stressors such as habitat loss (Chefaoui et al., 2018; Fourqurean et al., 2019; Taylor et al., 2026), or direct take (Carrasquero-Labarca et al., 2025; Humber et al., 2014; Lagueux et al., 2017) in particular areas (Table 3), may be important to arrest the current population trajectory in Tortuguero.

The adequate design, implementation and enforcement of MPAs is one of the main management tools used to mitigate these stressors in sensitive areas, conserving threatened species or restoring degraded habitats (Cramer et al., 2026b; Grorud-Colvert et al., 2021; Sanabria-Fernandez et al., 2019). Although MPAs can be highly effective as a conservation tool, monitoring and controlling these areas is still challenging (Davies et al., 2018; Wilhelm et al., 2014). This is particularly true in the Latin American and Caribbean region, where government priorities do not necessarily align with conservation outcomes and government control over distant regions is limited. Nicaragua is a good example in this context. In our study, 28 green turtles migrated from Tortuguero to two distinct foraging grounds in Nicaraguan waters. Most of these turtles went to Miskitos Keys where they remained completely and partially within the Marine Biological Reserve “*Cayos Miskitos*”.

Unfortunately this particular area (including the MPA) has traditionally been an area of large-scale legal extraction of green turtles, with up to an estimated of 12,000 adult or sub-adult turtles taken annually (Lagueux et al., 2014, 2017). This raises questions around the effectiveness of MPAs and the inclusion of financial mechanisms in their design to ensure their adequate implementation and effectiveness over long periods of time.

## 5. Conclusion

Regardless of a generalised recovery of green turtles worldwide and long-term conservation efforts, the nesting population at Tortuguero is showing a continuous decline in annual nest numbers which remains partially unexplained. Despite its global and regional importance, we are still learning about key aspects of movement ecology of green turtles nesting at Tortuguero. This study showcases conservation efforts for over 25 years of research, expanding our understanding of the migratory connectivity of nesting females, identifying their space use and protected area coverage, and elucidating the limitations of Tortuguero National Park and other MPAs in the Caribbean coast of Costa Rica to conserve the green turtle population nesting at this important rookery. We evaluated area use at 14 distinct locations, 11 of which were novel foraging grounds, and had never been assessed before. This provides us with a better understanding of green turtle migratory connectivity from Tortuguero, and insights into possible reasons for the current population decline that require further investigation.

## Acknowledgements

Constant monitoring and satellite tag deployments were conducted under annual research permits issued by Tortuguero Conservation Area from the National System of Conservation Areas of Costa Rica SINAC-ACTo. We thank park rangers and conservation authorities at Tortuguero National Park. The execution of this project was conducted under the approval of the University of Queensland’s Animal Ethics and Integrity Committee; Res. 2023/AE000357. The authors would like to thank and acknowledge the participation of dozens of research assistants and local staff from the Sea Turtle Conservancy monitoring program in Costa Rica; their effort and hard work contribute every year to the conservation of marine turtles and other species at this important rookery.

Funding for satellite transmitters was provided by the Rufford Foundation, Sea Turtle Conservancy, Shark Reef Aquarium, Pacsafe, TreadRight, Divinity Jewelry, SeaLife Trust, World Nomads, tarte Cosmetics, Greene Turtle Sports Bar & Grille, Pura Vida Bracelets, JD.com, honu Jewelry, Graft Cider, Gimme Snacks, Cinder & Salt, Certina, and Fahlo.

## Supplementary material

**Figure S1.**
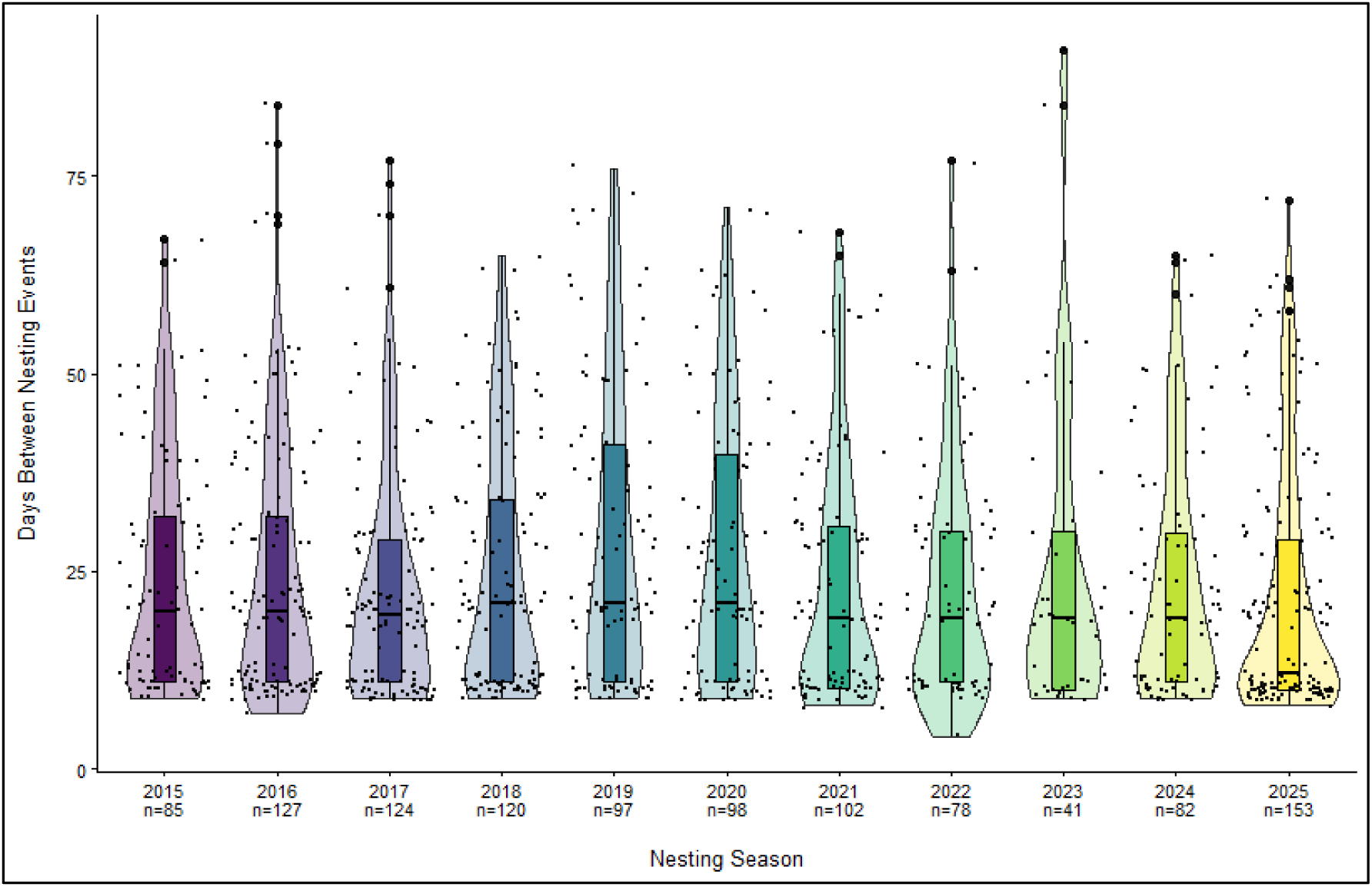
Annual internesting period for green turtles (*Chelonia mydas*) nesting at Tortuguero, Costa Rica, based on 10 years of mark-recapture information.

**Figure S2.**
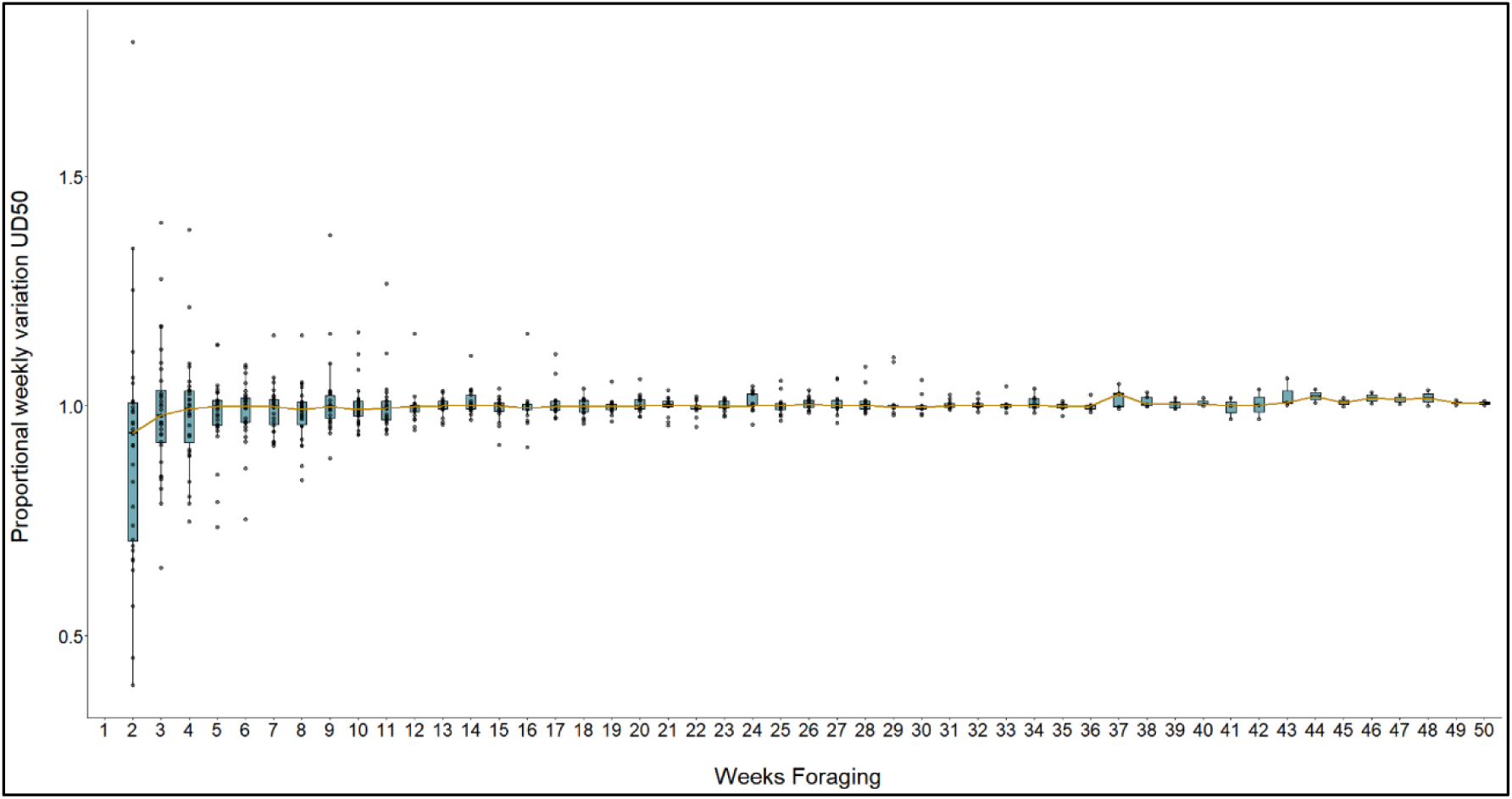
Proportional variation of weekly foraging UD50 for green turtles tracked from Tortuguero rookery.

**Table S.1.**
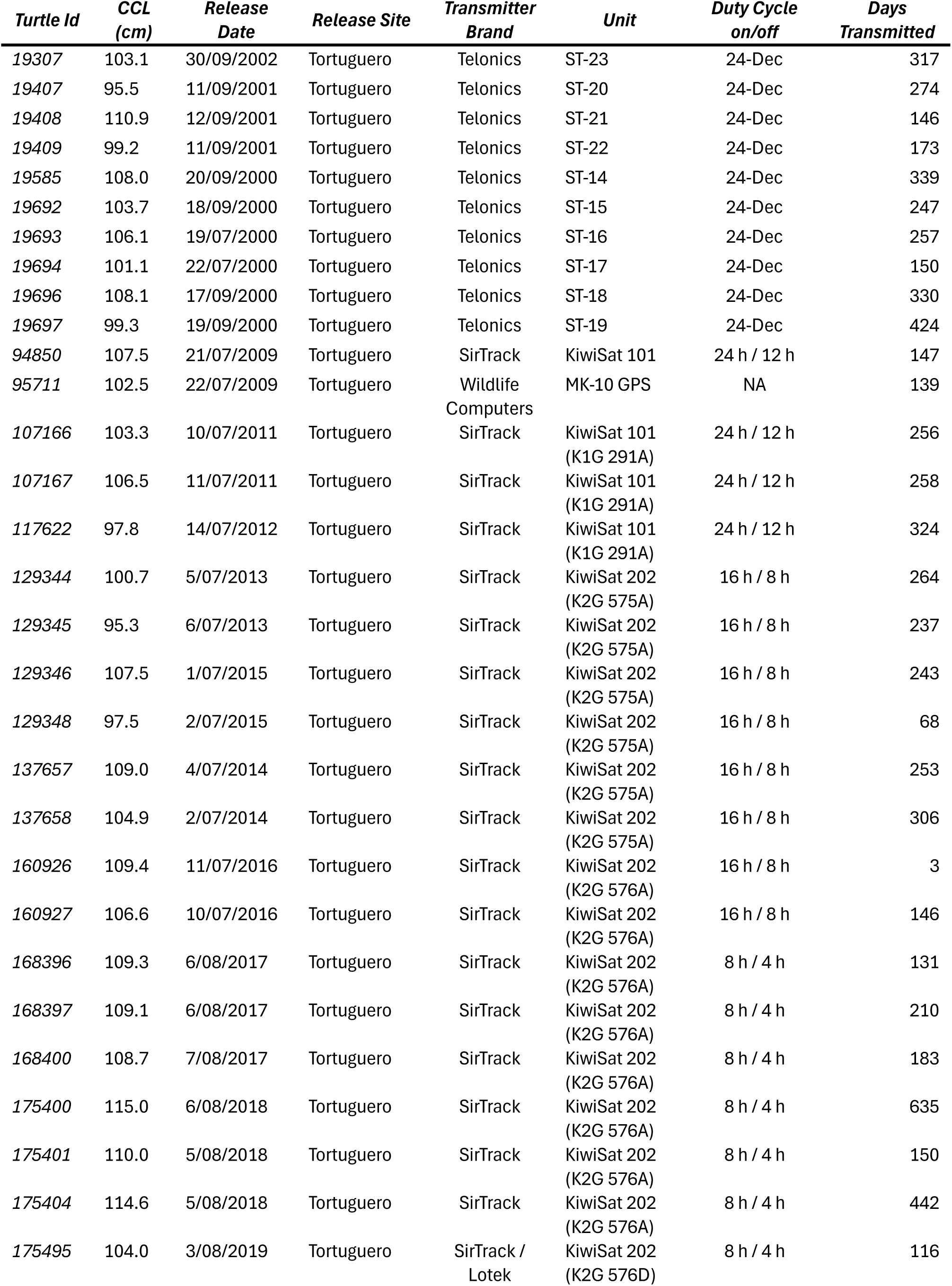
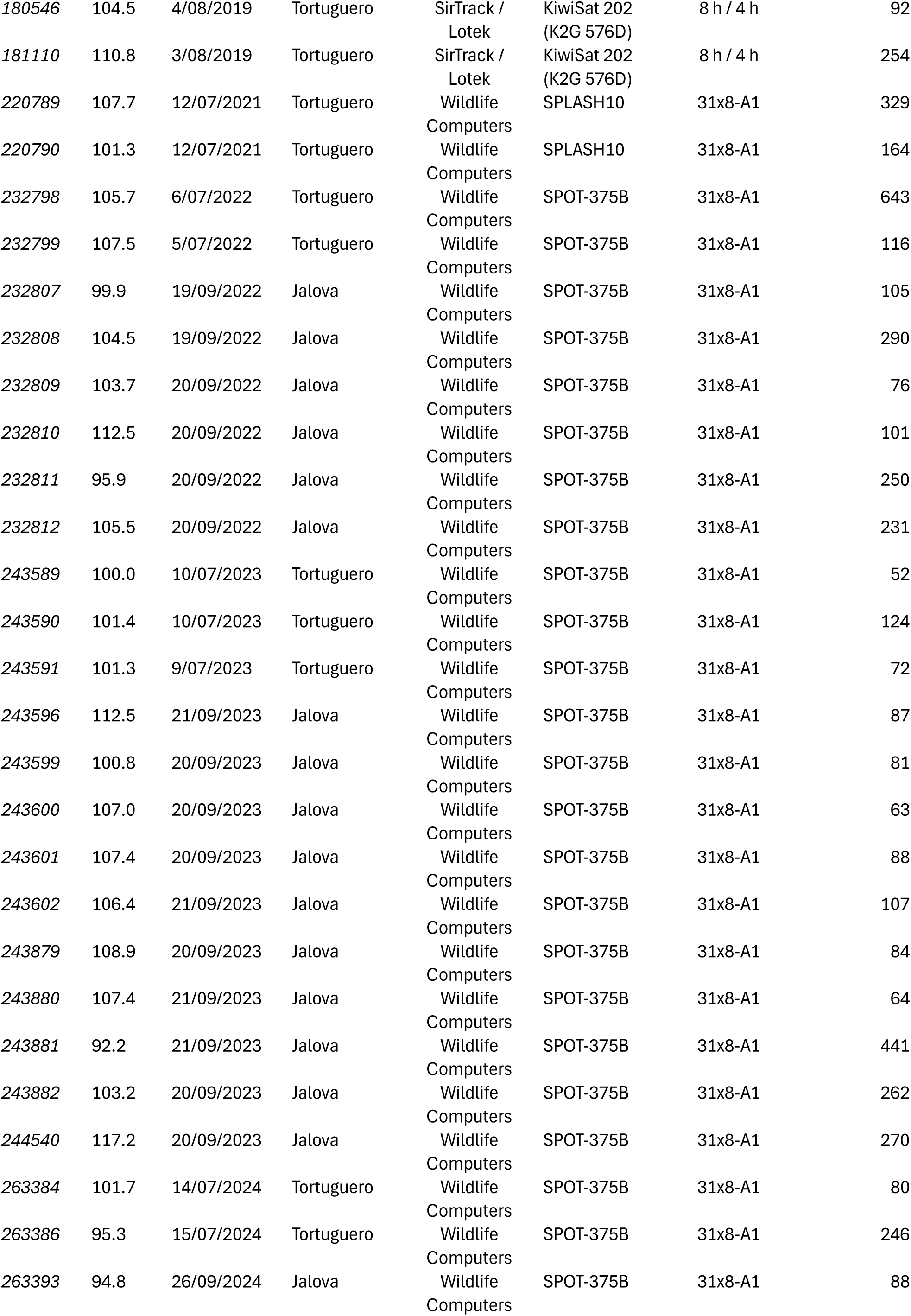
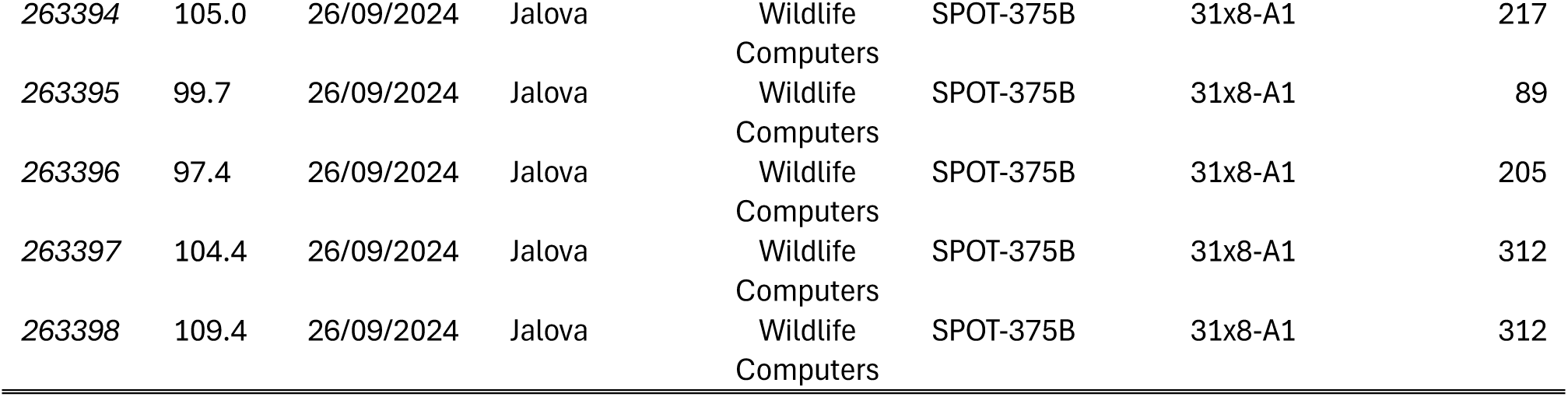
Satellite tag deployment information for green turtles (Chelonia mydas) nesting at Tortuguero, Costa Rica.

**Table S.2.**
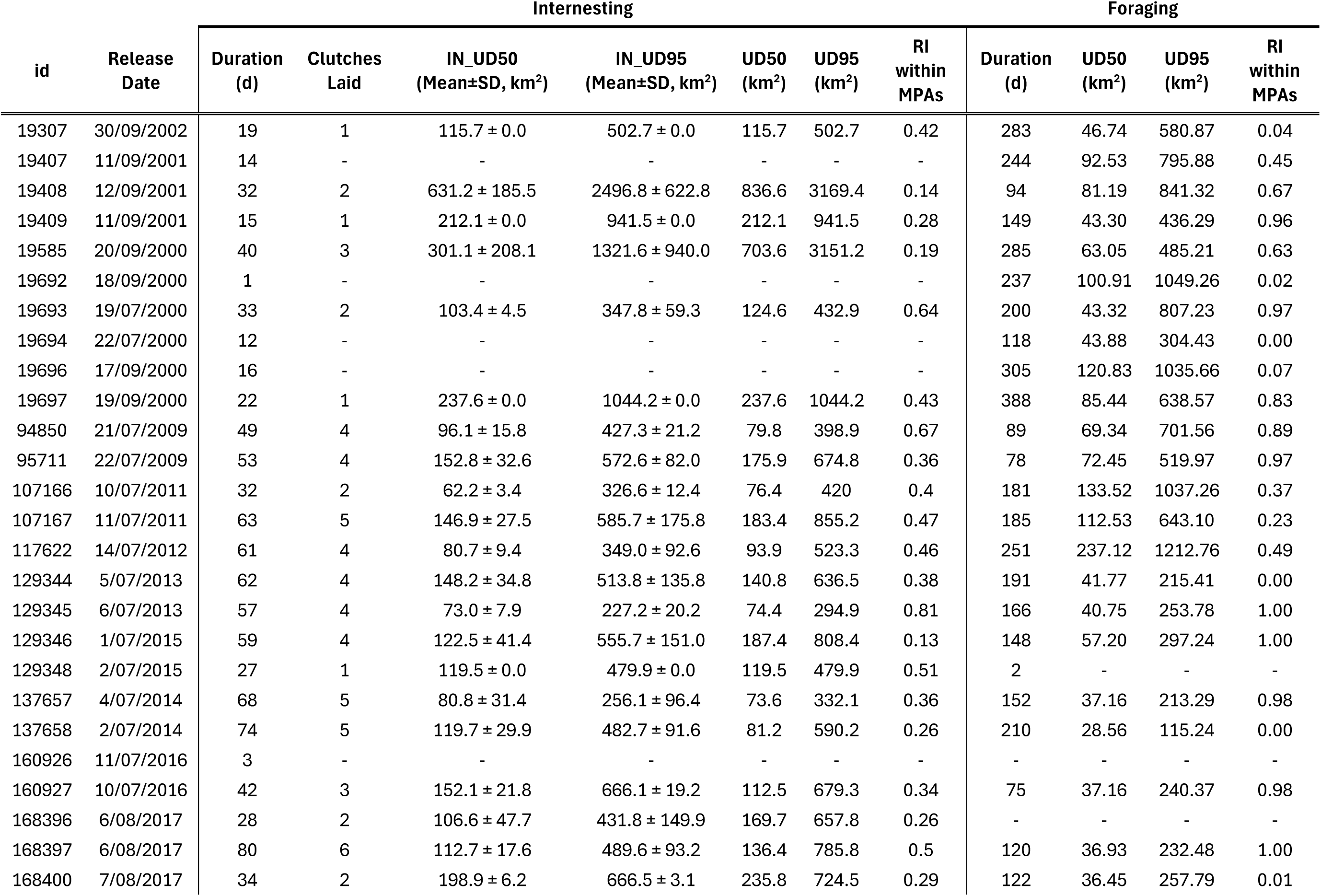
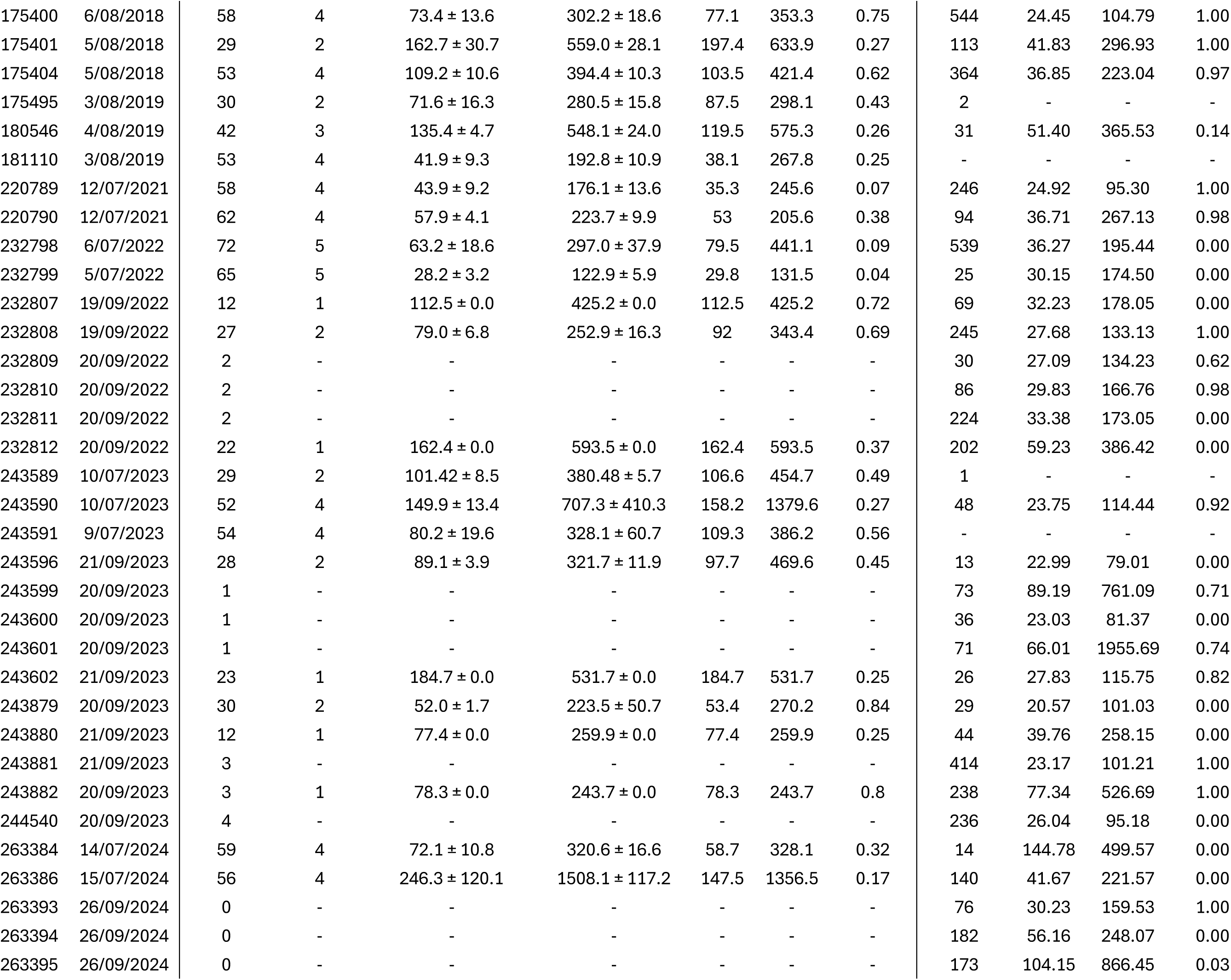
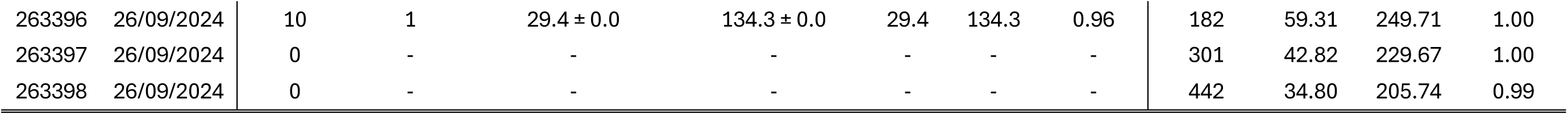
Internesting and foraging Utilization Distributions (UD) for tracked green turtles nesting at Tortuguero between 2000 and 2024. UD50 shows the core area used by each turtle during individual internesting and foraging activity. UD95 indicates the overall area used by each turtle during individual internesting and foraging activities. IN_UD50 and IN_UD95 are the mean core and overall areas in between nesting events for turtles that return to laid at least once after tag deployment RI within MPAs is the individual residency index, reflecting the proportion of time each individual spent within the boundaries of marine protected areas on either end of their migration cycle. The proportion of area in MPAs is the physical overlay of each UD95 with MPAs polygons (UNEP-WCMC 2026).

**Figure S3.**
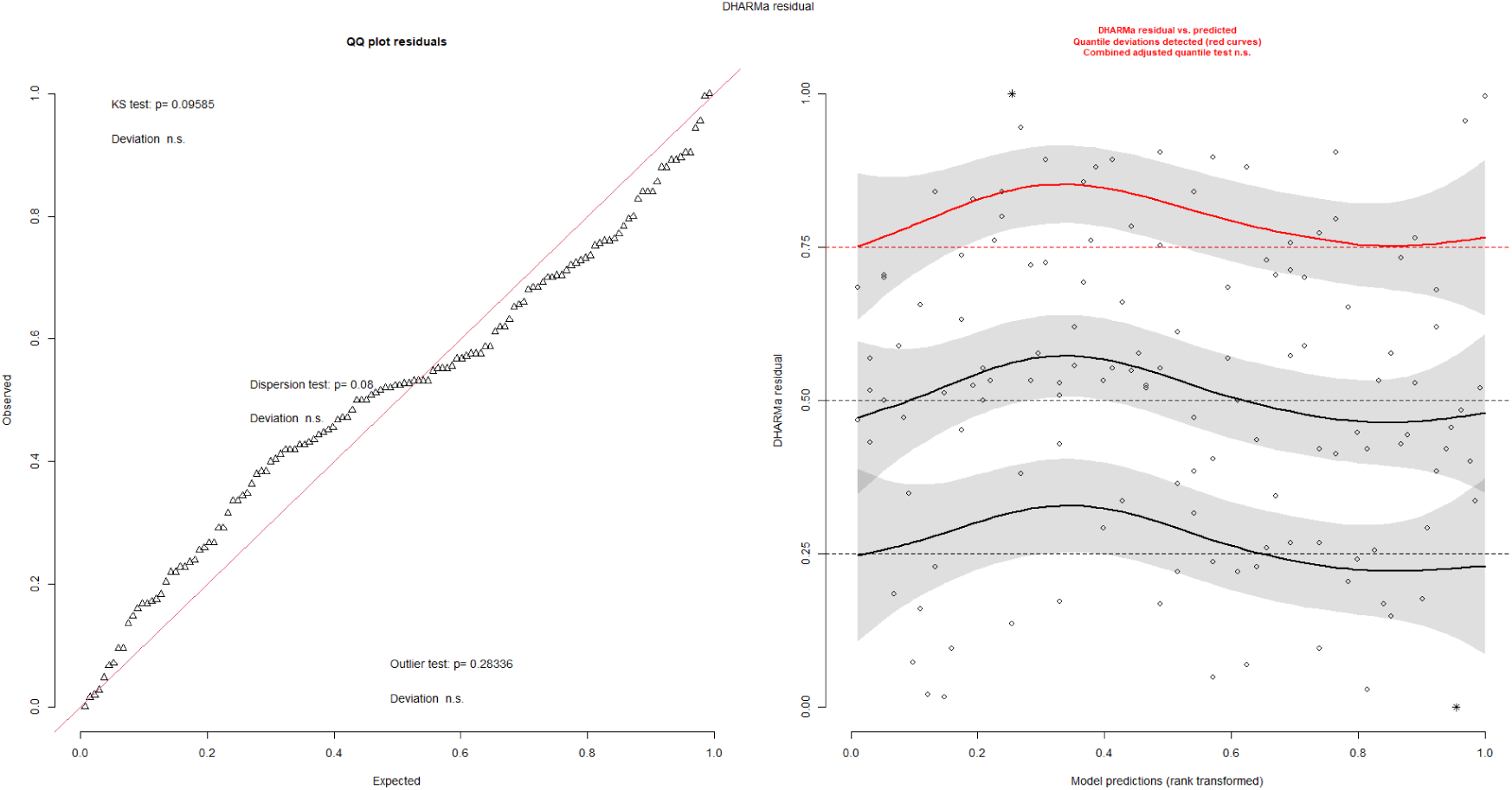
Diagnostic plots Internesting model 2. Evaluating UD50 over the internesting period for individual clutches and the effect of the release site.

**Figure S4.**
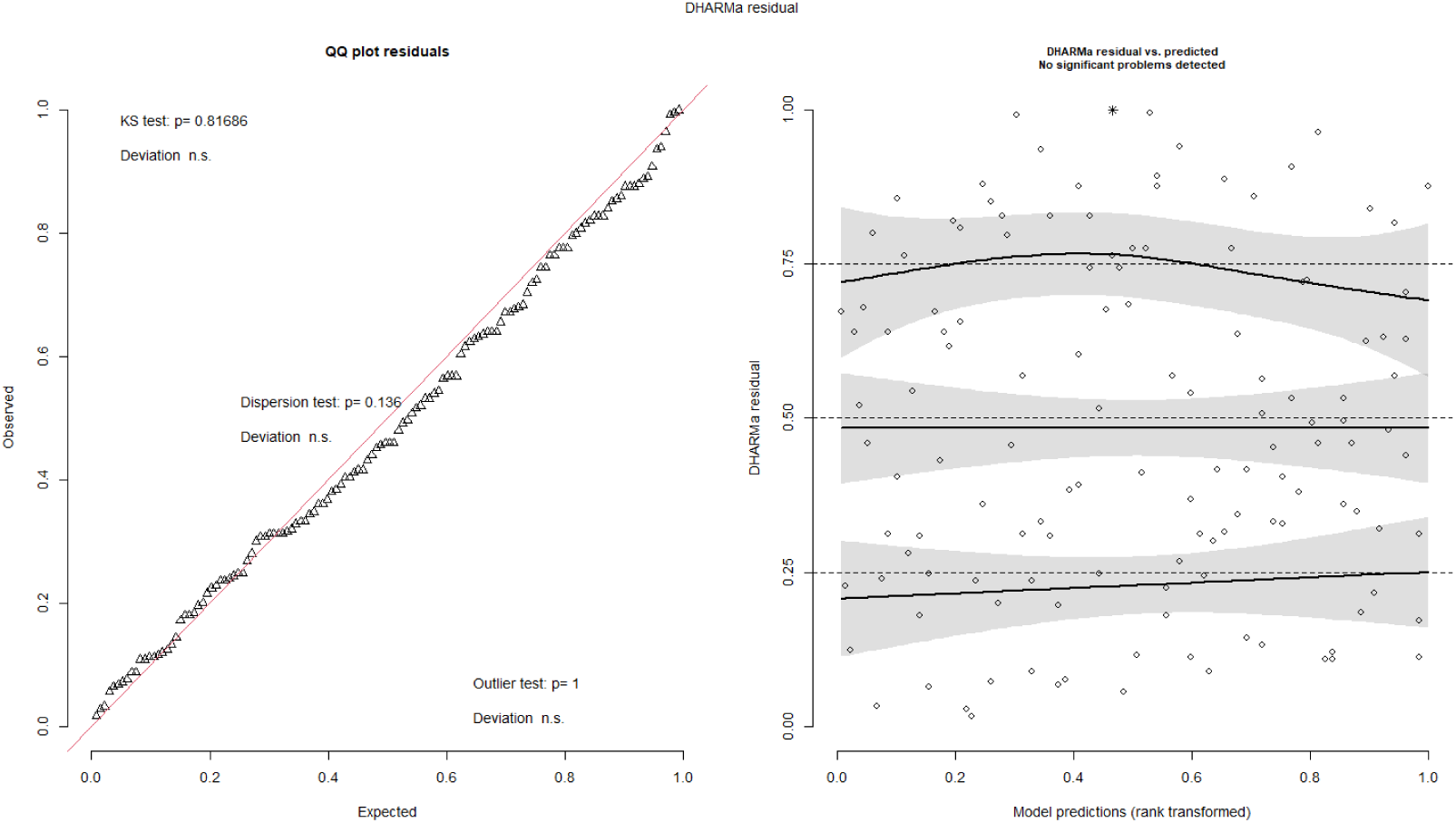
Diagnostic plots Internesting model 3. Evaluating the internesting residency index within MPAs over the internesting period for individual clutches and the effect of the release site.

**Figure S5.**
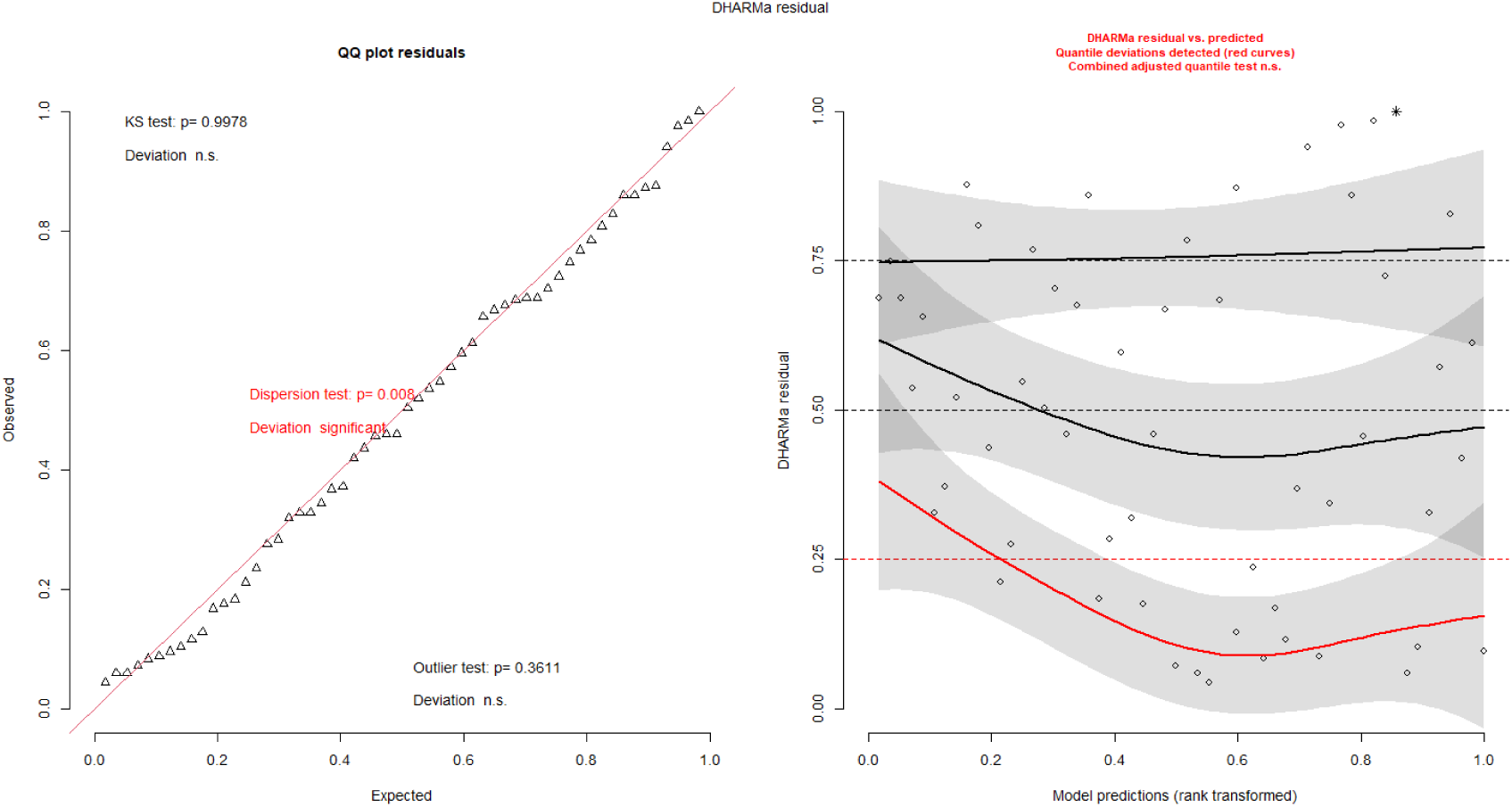
Diagnostic plots foraging model 1. Evaluating the foraging area (UD50) over time in every foraging ground.

**Figure S6.**
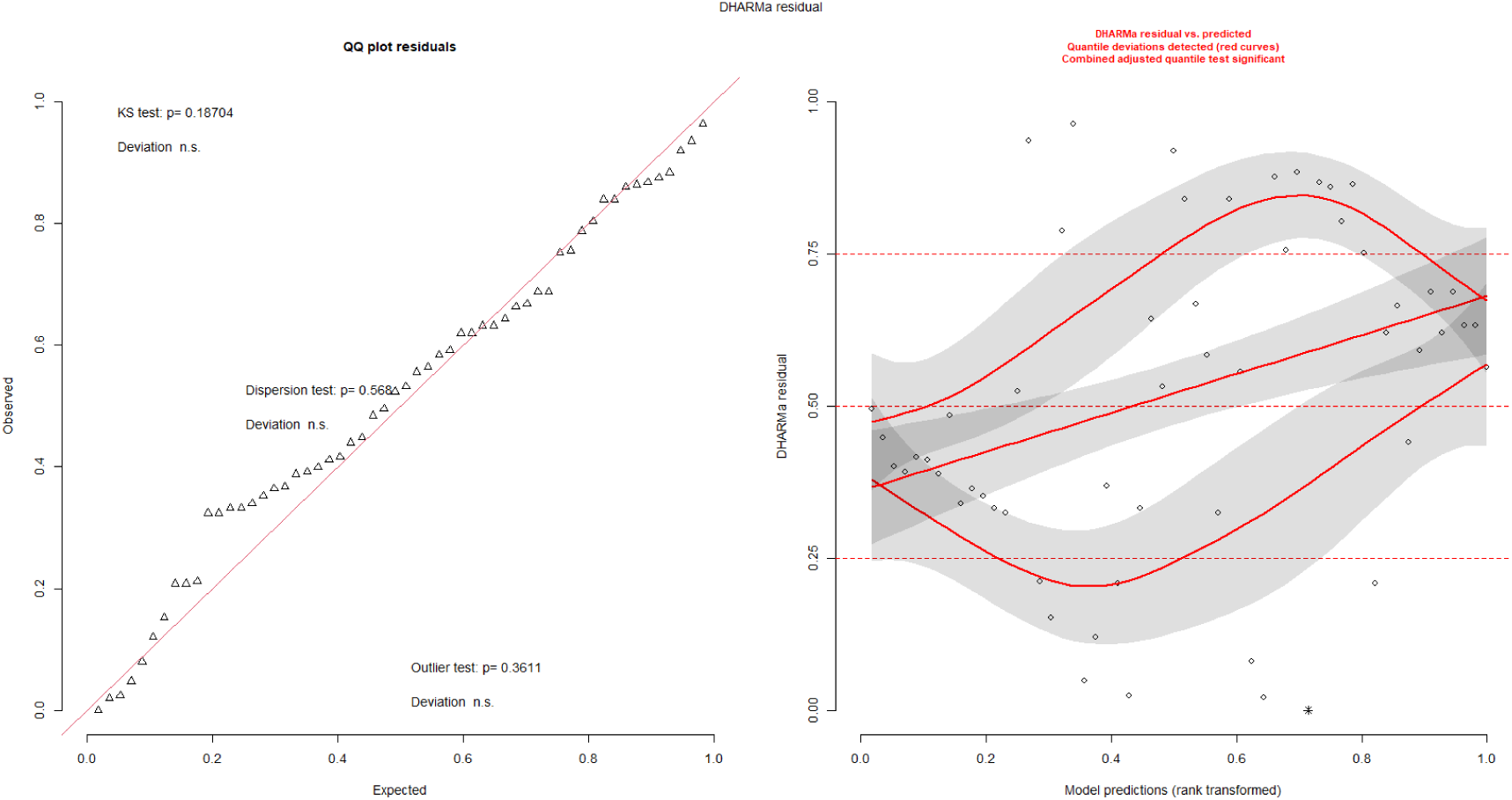
Diagnostic plots foraging model 2. Evaluating the residency index within MPAs compared to the individual foraging area (UD50) at each foraging ground.

**Table S.3.**
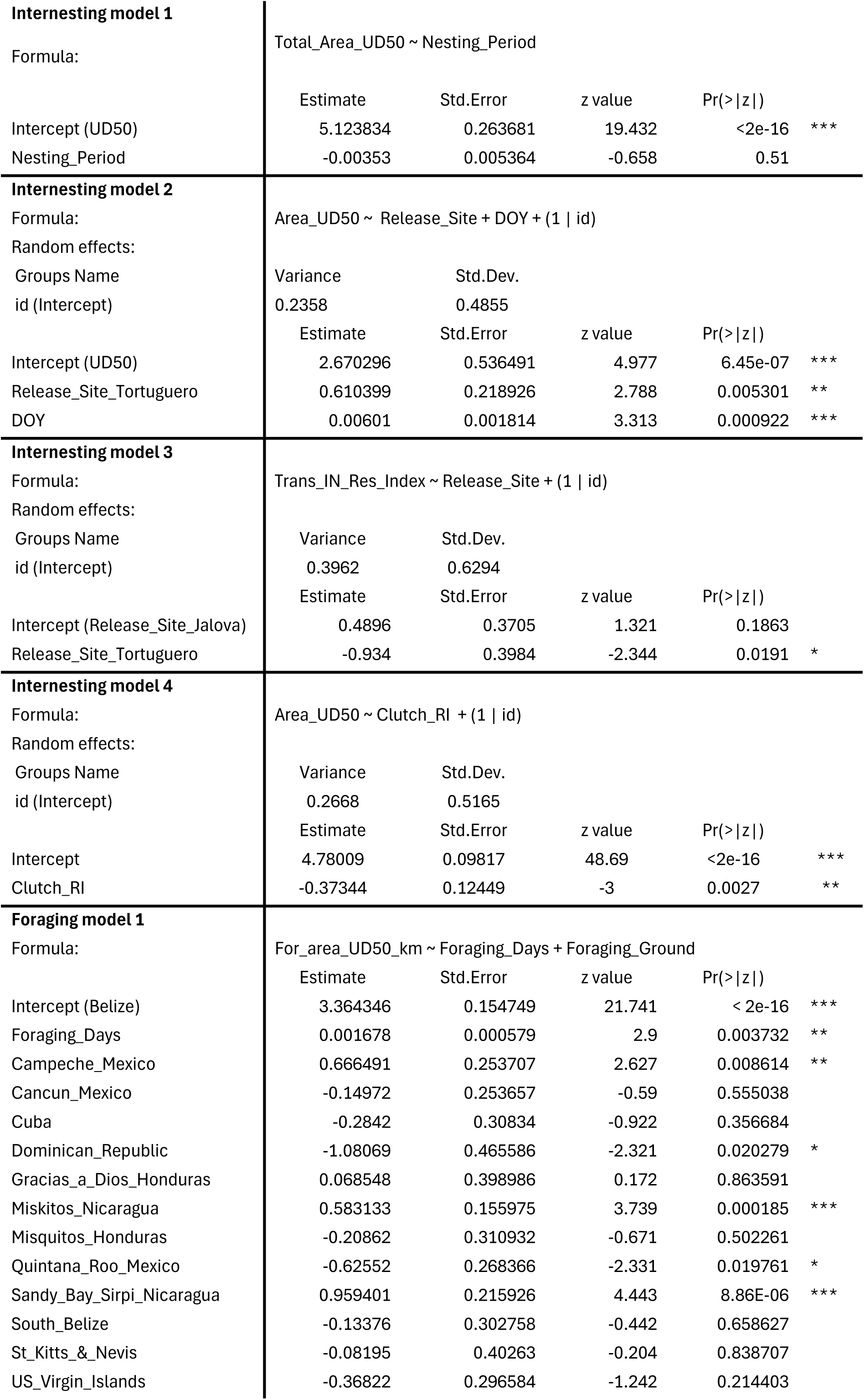
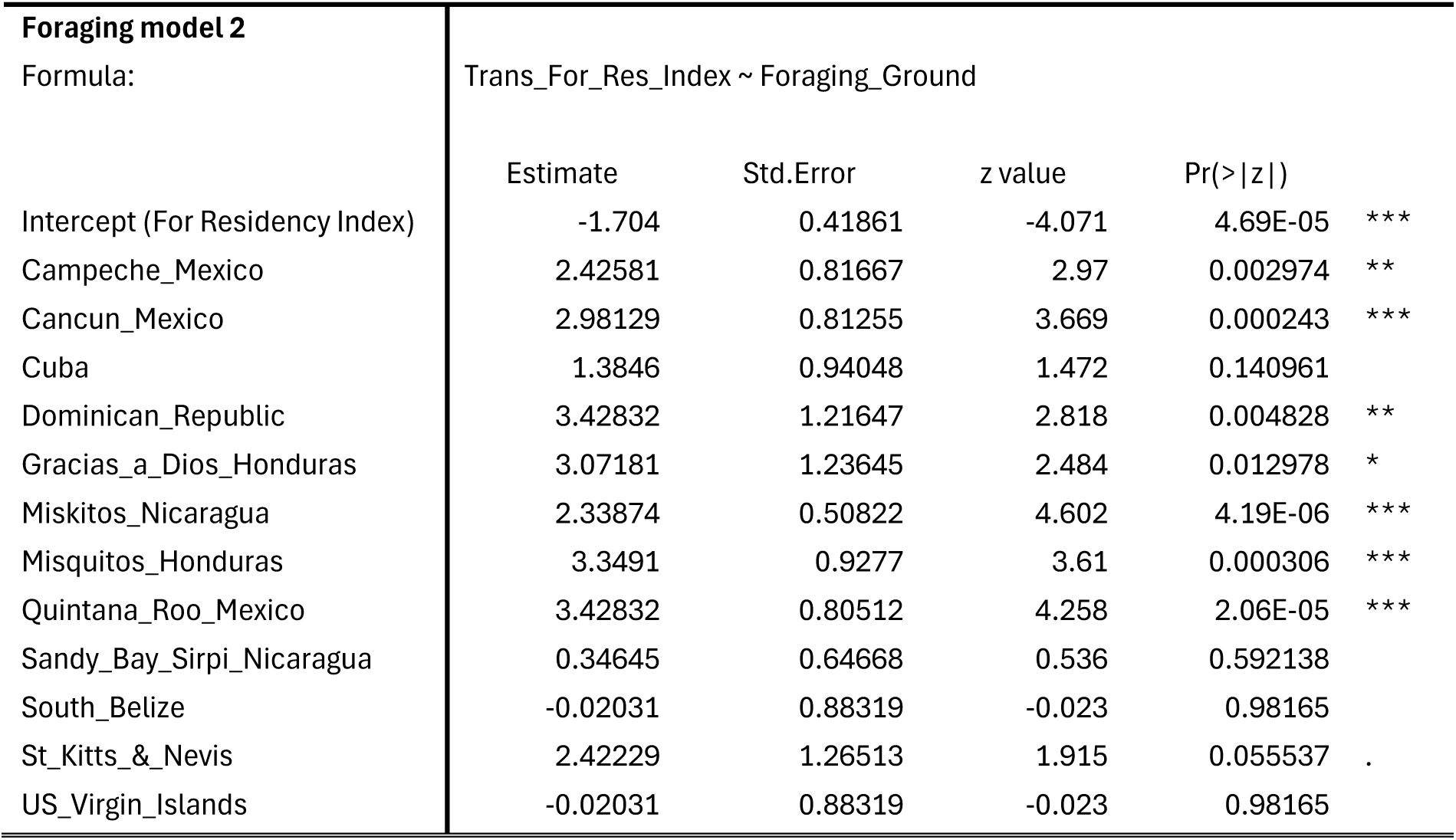
Outputs of generalised linear mixed models to assess the effect of variables on utilisation distributions (UDs) and residency index (RI) for green turtles during internesting and foraging behaviours.

